# Biophysical properties governing septin assembly

**DOI:** 10.1101/2021.03.22.436414

**Authors:** Benjamin L. Woods, Ian Seim, Jessica Liu, Grace McLaughlin, Kevin S. Cannon, Amy S. Gladfelter

## Abstract

Septin filaments build structures such as rings, lattices and gauzes that serve as platforms for localizing signaling and organizing cell membranes. How cells control the geometry of septin assemblies in poorly understood. We show here that septins are isodesmic polymers, in contrast to cooperative polymerization exhibited by F-actin and microtubules. We constructed a physical model to analyze and interpret how septin assemblies change in the presence of regulators in yeast extracts. Notably filaments differ in length and curvature in yeast extract compared to pure protein indicating cellular regulators modulate intrinsic biophysical features. Combining analysis of extracts from regulatory mutants with simulations, we found increased filament flexibility and reduced filament fragmentation promote assembly of septin rings, whereas reduced flexibility in crowded environments promotes local filament alignment. This work demonstrates how tuning of intrinsic features of septin filament assembly by regulatory proteins yields a diverse array of structures observed in cells.

## Introduction

Cells must bridge differences in scale when assembling and remodeling cytoskeletal structures that are micrometers in length out of nanometer-sized subunits. Microtubules and F-actin, two well studied cytoskeletal elements, are composed of dimeric and monomeric subunits, respectively, and form dynamic micron-scale assemblies essential for numerous cell processes. Similarly, septins, GTP-binding proteins conserved in many branches of eukaryotes, are a class of cytoskeletal-like proteins that form micron-scale higher-order assemblies from nanometer-scale building blocks (Auxier et al., 2019; Onishi and Pringle, 2016; Pan et al., 2007; Spiliotis and McMurray, 2020). Septin assemblies act as scaffolds for microtubules and F-actin, recruit essential proteins to the site of cytokinesis, and can organize the plasma membrane (Bridges et al., 2016; Clay et al., 2014; Gilden et al., 2012; Lew and Reed, 1995; Longtine et al., 2000; Spiliotis, 2010). These assemblies contribute to many cellular processes including cell division, cell migration, ciliogenesis, and neuronal branching (Dolat et al., 2014; Longtine et al., 1996; Mostowy and Cossart, 2012; Oh and Bi, 2011). Relevant to these varied cellular roles, septin dysfunction is linked to a variety of seemingly disparate human diseases, including multiple types of cancer and neuronal disorders (Angelis and Spiliotis, 2016; Montagna et al., 2015; Peterson and Petty, 2010).

The basic subunit of septin filament polymerization is a non-polar, hetero-oligomeric complex of septin proteins, ranging up to 32 nanometers long depending on the species (Bertin et al., 2008; Sirajuddin et al., 2007). Different organisms express different numbers of septin subunits ranging from 1 in the green algae *Chlamydomonas* to 13 different septin genes with many splice-variants and tissue-specific expression patterns in humans. Current work suggests that the functional state of septins is primarily in filaments and higher-order assemblies. Hetero-oligomeric septin complexes associate with the plasma membrane (and in some cells with other cytoskeletal elements), where they can anneal end-on with one another to polymerize into filaments (Bridges et al., 2014; McMurray et al., 2011; Smith et al., 2015; Spiliotis, 2018; Spiliotis et al., 2016). Septin filaments are then arranged into higher-order assemblies spanning microns such as rings, bundles, gauzes, and lattices (DeMay et al., 2009; Kinoshita et al., 2002; Ong et al., 2014). The mechanisms controlling septin filament polymerization and specifying the size and shape of higher-order assemblies remain poorly understood.

Cells control cytoskeletal polymerization in time and space to prevent constitutive self-assembly. Multiple described cytoskeletal polymers assemble cooperatively (**Fig 1A**) (Desai and Mitchison, 1997; Korn, 1982; Romberg et al., 2001). Cooperative polymers have two distinct phases in their assembly. First is an energetically unfavorable nucleation phase, typically requiring the association of more than two subunits. Interactions between individual monomers are weak, and multiple lateral associations between subunits stabilize a filament nucleus (Erickson, 1989; Oosawa and Kasai, 1962). After relatively slow nucleation, the elongation phase is characterized by rapid addition of subunits (**Fig 1B**). In the case of actin and microtubules, newly added subunits are stabilized via multiple interactions of the subunits at the end of the filament. As available monomers are incorporated into filaments, the available concentration of subunits decreases until the reaction reaches an equilibrium. In experiments with purified components that reach equilibrium, polymer lengths are observed to be normally distributed about an average length (**Fig 2A**), and unincorporated subunit concentration equals the critical concentration. Cooperative polymers are stabilized in the middle through the lateral associations between subunits and rarely fragment into separate filaments. Without severing regulatory proteins, cooperative polymers disassemble solely by subunits dissociating from the filament ends (**Fig 1A**). In cells, actin and microtubules length and arrangement are further controlled by regulators which nucleate, extend, cap, sever, or bundle filaments. Thus regulatory proteins enable assembly of the cytoskeleton into a variety of different higher-order structures (often with different filament properties and arrangements) enabling their diverse function (Pollard, 2016).

**Figure 1.**
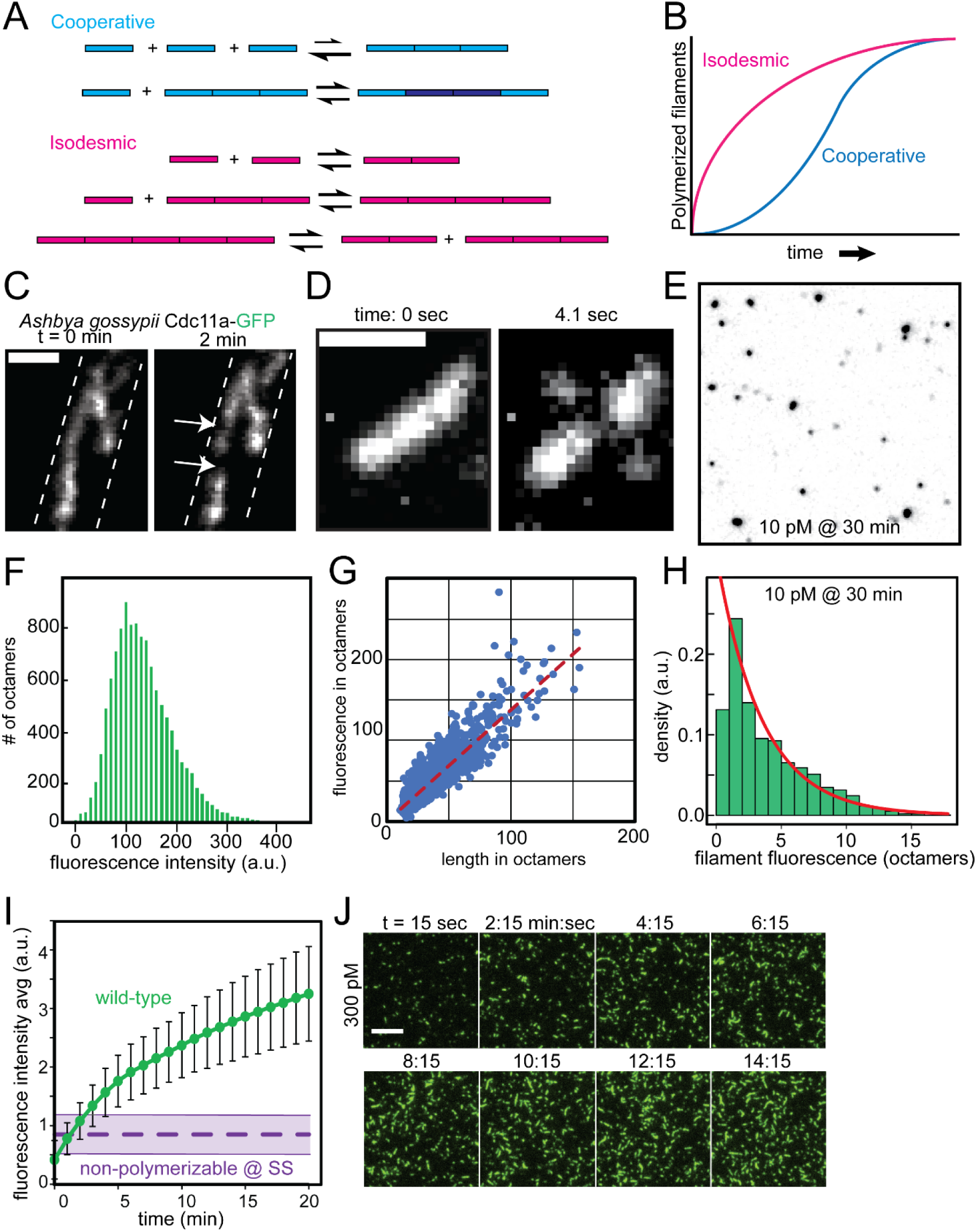
Evidence in support of isodesmic polymerization of septin filaments on membranes. A. Comparing features of cooperative (blue) and isodesmic (magenta) assembly. Cooperative polymers must first undergo energetically unfavorable nucleation prior to filament elongation at filament ends. Isodesmic polymers do not need nucleation prior to filament polymerization. Also, cooperative polymers disassemble primarily at filament ends, whereas isodesmic filaments can fragment along their length at the same rate as a subunit dissociating from the filament end. B. At low concentrations above a critical concentration, cooperative polymers (blue) must first undergo a slow nucleation phase prior to rapid filament elongation. Conversely, isodesmic polymers (magenta) do not have a nucleation phase, and polymerization steadily plateaus. C. Near-TIRF microscopy time lapse images of *Ashbya gossypii* expressing Cdc11a-GFP from the endogenous locus. Arrows demarcate sites of septin filament fragmentation. Dashed lines represent hyphal outlines. Scale bars, 2 μm. D. Recombinant septin filament polymerized on an SLB fragments at 4.1 sec. Scale bar, 1 μm. E. Septin filaments polymerized at 10 pM (final concentration) on SLBs after 10 min. Filaments are mostly fluorescent puncta of varying intensities. F. Histogram of fluorescence intensity of recombinant, non-polymerizable septin octamers harboring Cdc11α6-SNAP labeled with Alexafluor488 dye. The mode fluorescence of 110 a.u. was taken as that of a single septin octamer. G. Plot comparing measured filament length (in octamers) vs filament length estimated from fluorescence intensity (in octamers). H. Representative fluorescence distribution from an experiment as in **D**, where fluorescence of puncta is given as a multiple of the single octamer. The red line is an exponential fit. The plot represents measurements from > 5,900 filaments. I. Averaged fluorescence intensity (per pixel) of wild-type recombinant septin filaments over time (reaction concentration: 200 pM, n = 3; error bars represent standard error of the mean) assessed by total fluorescence over background of a 7744 µm^2^ patch. This is compared to the total fluorescence of septins capped the non-polymerizable mutant Cdc11α6 (reaction concentration: 200 pM, n = 4). Dashed line represents average Cdc11α6 intensity, area represents standard error of the mean. J. Recombinant septin octamers (300 pM, with Cdc11-SNAP fluorescently labeled with Alexafluor488 dye) incubated on SLBs polymerize into filaments over time as visualized by TIRF microscopy. Scale bar, 5 μm

**Figure 2.**
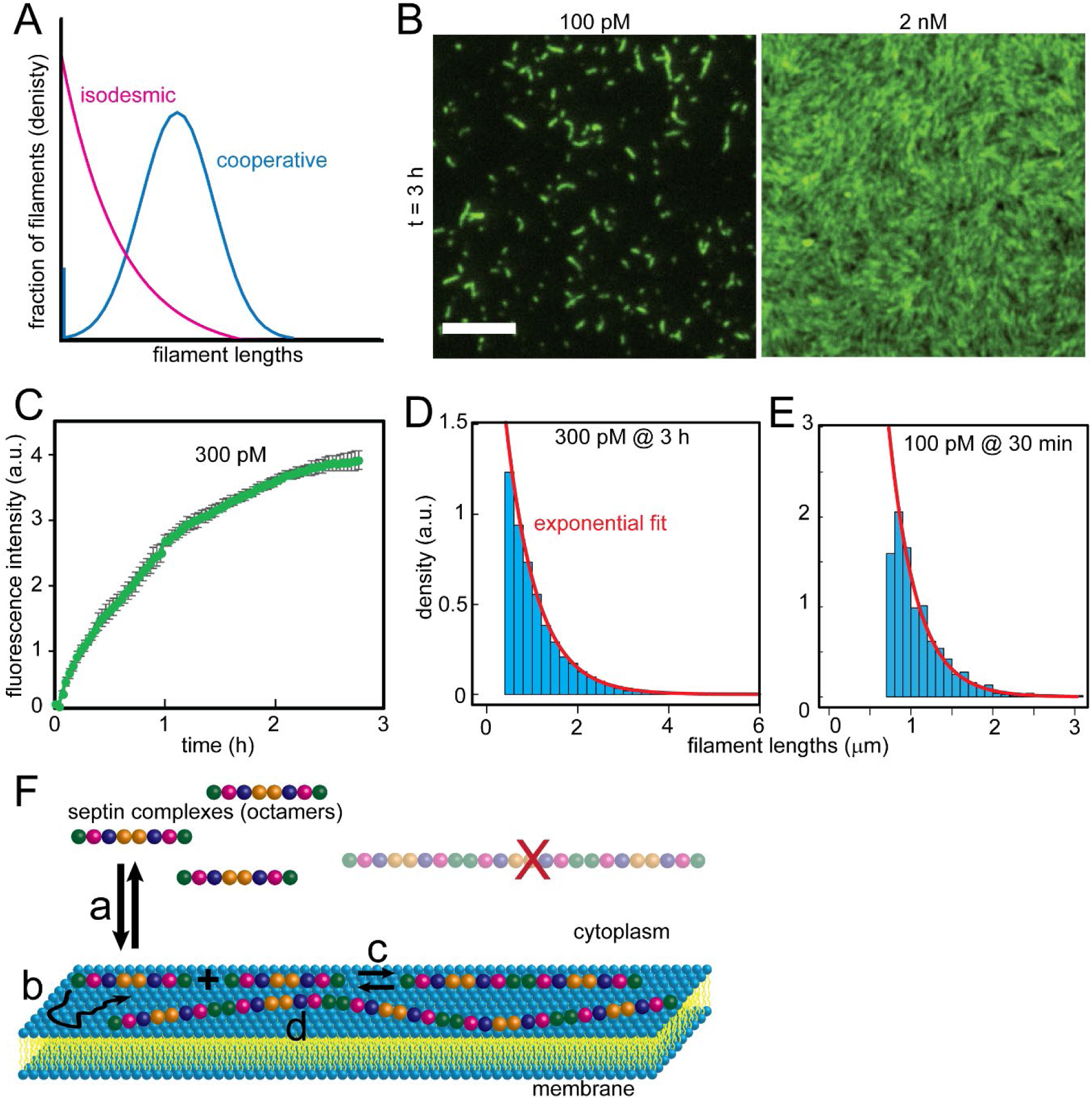
Septin filaments lengths are exponentially distributed. **A.** Predicted filament length distributions of cooperative (purple) and isodesmic (magenta) polymers at steady state. **B.** Septins at indicated reaction concentrations were incubated on planar supported phospholipid bilayers and allowed to polymerize for >three hours. Filaments were imaged by TIRF microscopy. Scale bar, 5 μm. **C.** Septin adsorbance to the membrane as assessed by fluorescence intensity above background over time (n = 3). Error bars represent standard error of the mean. **D.** Representative filament length distribution histogram at steady state, with a fitted exponential (red line). **E.** Representative filament length distribution before reaching steady state, with a fitted exponential. **F.** Cartoon diagram of septin filament assembly reactions: **a.** septin binding/unbinding to membrane, **b.** septin diffusivity at the membrane, **c.** septin-septin annealing/fragmenting (i.e. polymerization), and **d.** filament flexibility. Polymerized filaments do not appreciably unbind the membrane.

In comparison to actin and microtubules, very little is known about septin filament polymerization. Recent biochemical analysis indicates that bulk adsorption of septins onto curved membranes is a cooperative process due to the collective membrane-avidity of septin complexes incorporated into filaments (Cannon et al., 2019; Cannon et al., 2017). As septin complexes polymerize into filaments, the septin dissociation rate from the membrane decreases (**Fig 2F**). However, those studies measured membrane *adsorption,* which is distinct from filament *polymerization*. Previous work developed a minimal, reconstituted septin filament polymerization assay *in vitro*, incubating recombinantly-expressed septin complexes from the budding yeast *S. cerevisiae* on planar, supported phospholipid bilayers (Bridges et al., 2014). Canonical budding yeast septin complexes are hetero-octamers (Shs1/Cdc11-Cdc12-Cdc3-Cdc10-Cdc10-Cdc3-Cdc12-Cdc11/Shs1). When incubated on SLBs, septin octamers capped with Cdc11 bound the membrane where they diffuse, collide, and anneal end-on with one another to polymerize into larger, flexible filaments. Notably, septin filaments were observed to fragment *without* the addition of specific severing proteins, uncharacteristic of a cooperative polymer (Bridges et al., 2014). This raises the possibility that septin filament polymerization is not cooperative, but instead an isodesmic process.

Isodesmic polymers do not have separate nucleation and elongation phases (Romberg et al., 2001). This is because isodesmic polymers have identical bonds at each step of polymerization, meaning that any two subunits have the same affinity for each other irrespective of their position within the filament and thus fragment at a frequency proportional to their length (**Fig 1A**). Thus, isodesmic filament lengths are exponentially distributed with a greater number of short filaments than long filaments at equilibrium (**Fig 2A**). There are few examples of isodesmic cytoskeletal elements in biology. The classic example is the polymerization of glutamate dehydrogenase, which occurs with kinetics typical of protein-protein association (k_A_= 1.5 µM^-1^s^-1^ and k_D_ = 5 s^-1^) (Thusius et al., 1975). A more recent study has modeled assembly of actin from the eukaryotic parasite *Toxoplasma gondii* as isodesmic; however the kinetics are several orders of magnitude slower than typical protein-protein associations (Skillman et al., 2013).

How cells dynamically build and shape higher-order septin assemblies remains an ongoing area of intensive study (Beber et al., 2019; Bertin et al., 2010; Bridges et al., 2016; Bridges et al., 2014; DeMay et al., 2011; DeMay et al., 2009; McDonald et al., 2017; Okada et al., 2013; Ong et al., 2014; Sadian et al., 2013). Genetic and biochemical studies in multiple model systems have identified a network of septin regulators, including kinases, cell polarity proteins, and the cell cycle machinery (Bi and Park, 2012; Oh and Bi, 2011). To understand this regulatory network, it is important to study the intrinsic properties of septin polymers. Here we build upon the previously published filament polymerization reconstitution assay where purified septins are assembled into filaments on two-dimensional planar SLBs (Bridges et al., 2014). We develop a molecular dynamics model of isodesmic filament polymerization in two-dimensions based on measured parameters. Finally, we use whole cell extracts in the reconstitution assay to begin dissecting the role of septin regulators in building different features of higher-order assemblies.

## Results

We set out to determine whether septin polymerization occurs in a cooperative or isodesmic manner on membrane surfaces (**Fig 1A**). We previously observed that septin filaments in *Schizosaccharomyces pombe* and the filamentous fungus *Ashbya gossypii* cells fragment, which suggests an isodesmic assembly (Bridges et al., 2014) (**Fig 1C**). Septin filaments formed on SLBs with recombinant protein also readily fragment indicating that cellular factors such as severing proteins are not essential for this process (Bridges et al., 2014; Khan et al., 2018) (**Fig 1D**). Here, we report two additional lines of evidence that suggest septin polymerization on membranes is isodesmic: 1) no critical concentration and 2) exponentially-distributed filament lengths.

### Septins polymerize into filaments without a detectable critical concentration

Cooperative polymerization requires a minimal (critical) subunit concentration to form stable nuclei (**Fig 1A**). In contrast, isodesmic polymerization does not require nucleation and there is no minimal concentration necessary to form polymers. We tested whether there is a critical concentration necessary for septin filament polymerization on SLBs. Fluorescently-tagged, recombinantly-expressed *S. cerevisiae* septins were adsorbed onto planar SLBs from a range of solution concentrations down to 10 pM. At very low concentrations, septins visualized were mostly diffraction-limited puncta of varying brightness (**Fig 1E**). Based on our previous work, we assume that each punctum is a filament of various length (Bridges et al., 2014; Cannon et al., 2019). To estimate the number of septin complexes in puncta we first determined the fluorescence of single non-polymerizable septin octamers (**Fig 1F**). Given the propensity of septins to form paired filaments, we then needed to determine if paired or unpaired filaments form with recombinant proteins on SLBs to accurately interpret the fluorescence intensity of puncta. We measured the fluorescence of longer, resolvable filaments and found fluorescence correlated with the number of octamers predicted by length, suggesting filaments recombinant septins polymerize into unpaired, single filaments (**Fig 1G**). We then estimated the number of octamers in each punctum as the closest integer multiple of the single octamer. Even at the lowest concentration tested (10 pM), we detected assemblies that are consistent with filaments up to 15 octamers in length (480nm) (**Fig 1H**). Thus, we were unable to detect a concentration insufficient for polymerization on SLBs. Evidence of detectable filaments at very low concentrations that are orders of magnitude lower than cellular concentrations (100-200 nM) suggests that there is not a critical concentration for septin polymerization (Bridges et al., 2014).

Even at these very low concentrations, we observed that filaments formed relatively quickly. We assume that single octamers are reversibly bound to the SLB, and an equilibrium is quickly established between SLB-bound and solution octamers. As septin octamers polymerize into filaments their binding to the SLB will become irreversible. This likely happens at the level of a dimer of octamers, since a cooperative association that doubles the bond interface increases the association equilibrium constant by many orders of magnitude (Erickson, 1989). The concentration of reversibly bound monomeric octamers should remain constant, so these irreversibly bound filaments will cause a progressive increase in the total bound septins. At somewhat higher concentrations (200 – 300 pM), there was a clear, progressive increase in fluorescence and detectable filaments for wild-type complex but not a non-polymerizable complex (**Fig 1I & J**). This was consistent with what had been seen previously at > 5X higher protein concentrations (Bridges et al., 2014). These conditions led to clearly resolved filaments up to 1.0 μm long (∼30 octamers). We were unable to detect a kinetic lag in filament polymerization and septin adsorption on planar SLBs.

### Septin filament lengths are exponentially distributed

Another feature distinguishing cooperative and isodesmic polymers is their filament length distributions at steady state. Once polymerization reaches steady state (i.e. when rates of polymerization and depolymerization are equal) lengths of cooperative polymer are distributed normally about an average length with a subpopulation of free monomers that equals the critical concentration (**Fig 2A**). In contrast, isodesmic polymers fragment at a rate proportional to their length, meaning at steady state isodesmic polymer lengths are exponentially distributed with more short filaments than long filaments. To determine the septin filament length distribution we measured the filament lengths after incubating septins on SLBs for several hours until a steady state was reached when bulk adsorption (as determined by fluorescence) plateaued (**Fig 2B & C**). At concentrations above 500 pM, septin filaments were too densely packed on SLBs at steady state to individually resolve and measure their lengths (**Fig 2B**). However, below this concentration, lengths were clearly exponentially distributed both at steady state and prior to reaching steady state (**Fig 2D, E**), consistent with polymerization being isodesmic. Similarly, as shown above, at very low concentrations (10 pM), length distributions were also exponentially distributed as assessed by fluorescence (**Fig 1H**).

Collectively, 1) evidence of filament fragmentation, 2) the absence of a critical concentration, and 3) the exponential distribution of septin filament lengths suggest that septin filament polymerization on membranes is isodesmic. How does isodesmic polymerization of septins on membranes impact assembly of higher-order structures seen in cells? Without well characterized examples of isodesmic polymers in biology, it is unclear how cells control septin filament assembly. We hypothesize that like cooperative polymers, regulators tune specific biophysical reactions of isodesmic polymerization to build distinct higher-order structures. Although there are numerous proposed septin regulators from genetic and biochemical studies, how they regulate isodesmic polymerization is mysterious.

### Parameterizing a physical model of isodesmic septin filament assembly

To begin to examine how different regulatory mechanisms could produce distinct septin assemblies, we constructed a molecular dynamics model of isodesmic polymer assembly on membranes. The model is constructed from a hypothesized minimal set of biophysical reactions necessary to simulate filament polymerization on a supported lipid bilayer (**Fig 2F**). In the model we consider septin filament polymerization as the product of several, sequential reactions: **a**) binding (and unbinding) of septin octamers to the membrane, **b**) octamer diffusion at the membrane, and **c**) octamer-octamer end-on interactions (annealing and filament fragmentation) at the membrane. We model septin octamers with excluded volume interactions, enabling us to capture crowding and steric effects on filament polymerization while explicitly encoding **d**) filament flexibility which we hypothesize also influences the organization of higher-order assemblies. In order to parameterize the physical model, we measured biophysical reactions from the *in vitro* reconstitution assay using recombinant septins.

### Measuring septin octamer-membrane binding affinity on planar supported phospholipid bilayers

We first determined the septin octamer binding affinity to membranes. We incubated fluorescently tagged, recombinant cdc11α6-capped septin octamers on supported phospholipid bilayers and measured septin-membrane binding and unbinding events by TIRF microscopy. The number of binding events were counted as a function of the septin concentration over time per observable binding area at steady state. The association rate on planar SLBs was **2.11 ± 0.24 μm^-2^ s^-1^ nM^-1^** (**Table 1**). Next, to determine the septin-membrane disassociation rate, septin octamer dwell times were measured and data was fit to an exponential curve. The dissociation rate calculated from dwell times for single octamers on planar SLBs was **1.42 ± 0.22 s^-1^** (**Fig 3A; Table 1**). The octamer-membrane affinity reported here is higher than what was previously reported on curved membranes (Cannon et al., 2019), however this likely reflects differences in the ionic strength between these experiments.

**Figure 3.**
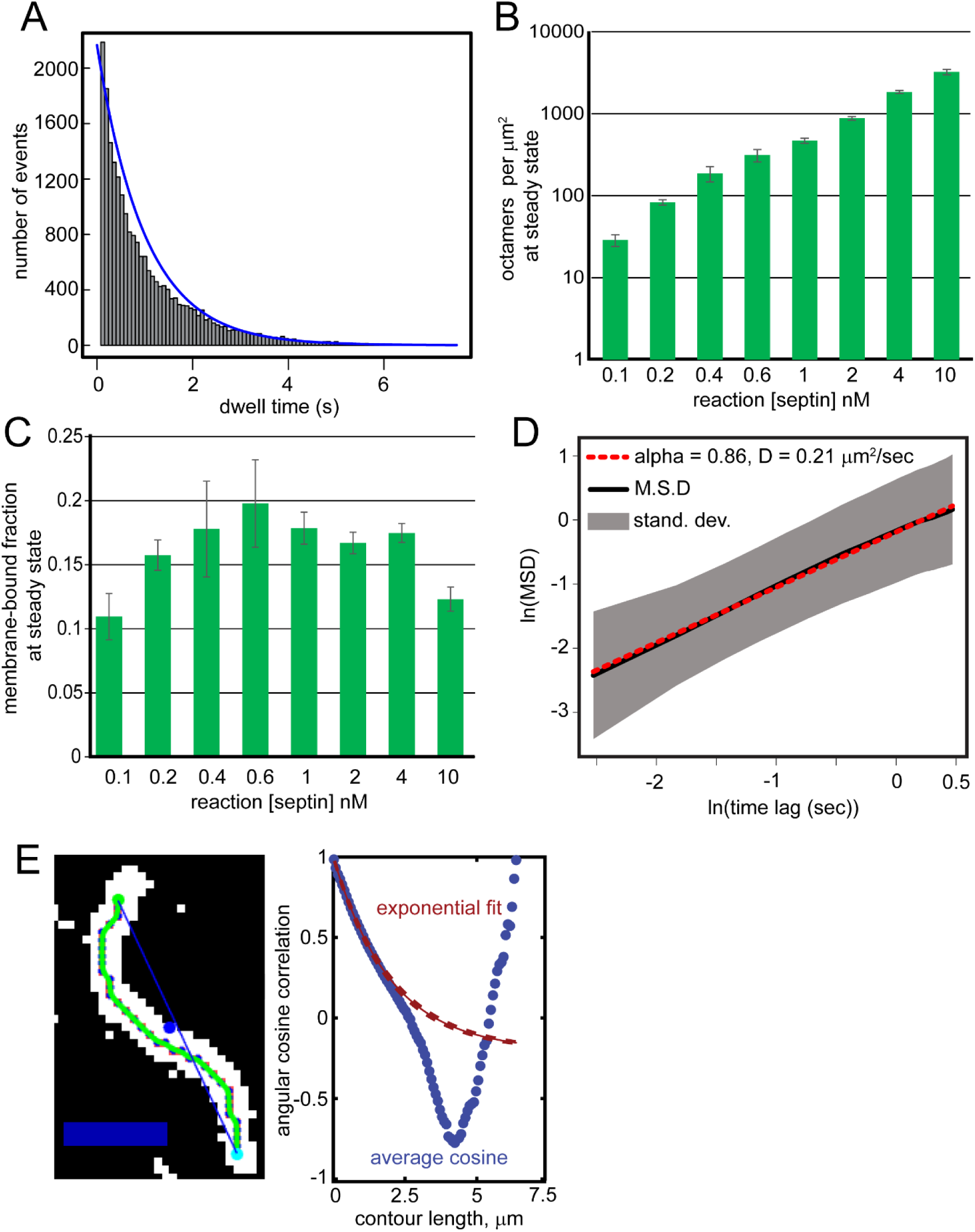
Biophysical properties of septins during assembly. A. A representative dwell time histogram of non-polymerizable septin octamers (>23,000 events) incubated on planar supported phospholipid bilayers. B. Octamer density measurements assessed by adsorption (bulk fluorescence) via TIRF microscopy and indicated reaction concentrations (averages from n = > 5 TIRF images from 3-5 separate experiments at each concentration). C. Values from **B**, estimating the membrane-bound fraction of the total reaction septin concentration. The average membrane density is extrapolated over entire SLB. D. Means squared displacement (M.S.D) plot averaged from three experiments with a fit of the octamer diffusion on planar supported phospholipid bilayers. Red dashed line is the diffusion fit. Black is the mean. Gray region is the standard deviation of the average M.S.D. E. (Left) Example of the filament contour length (green line) relative to its Fourier length (blue line connecting filament ends). Scale bar, 1 μm. (Right) Plot describing the cosine correlation as a function of contour length. For each filament, tangent angles between points on the filament are measured along its contour length. The average cosine of these tangent angles is plotted as a function of the contour length. The persistence length (**L_P_**) is extracted from an exponential fit to the plotted average cosine correlation. Measurements are from 1073 filaments averaging 1.62 μm in length.

**Table 1.**
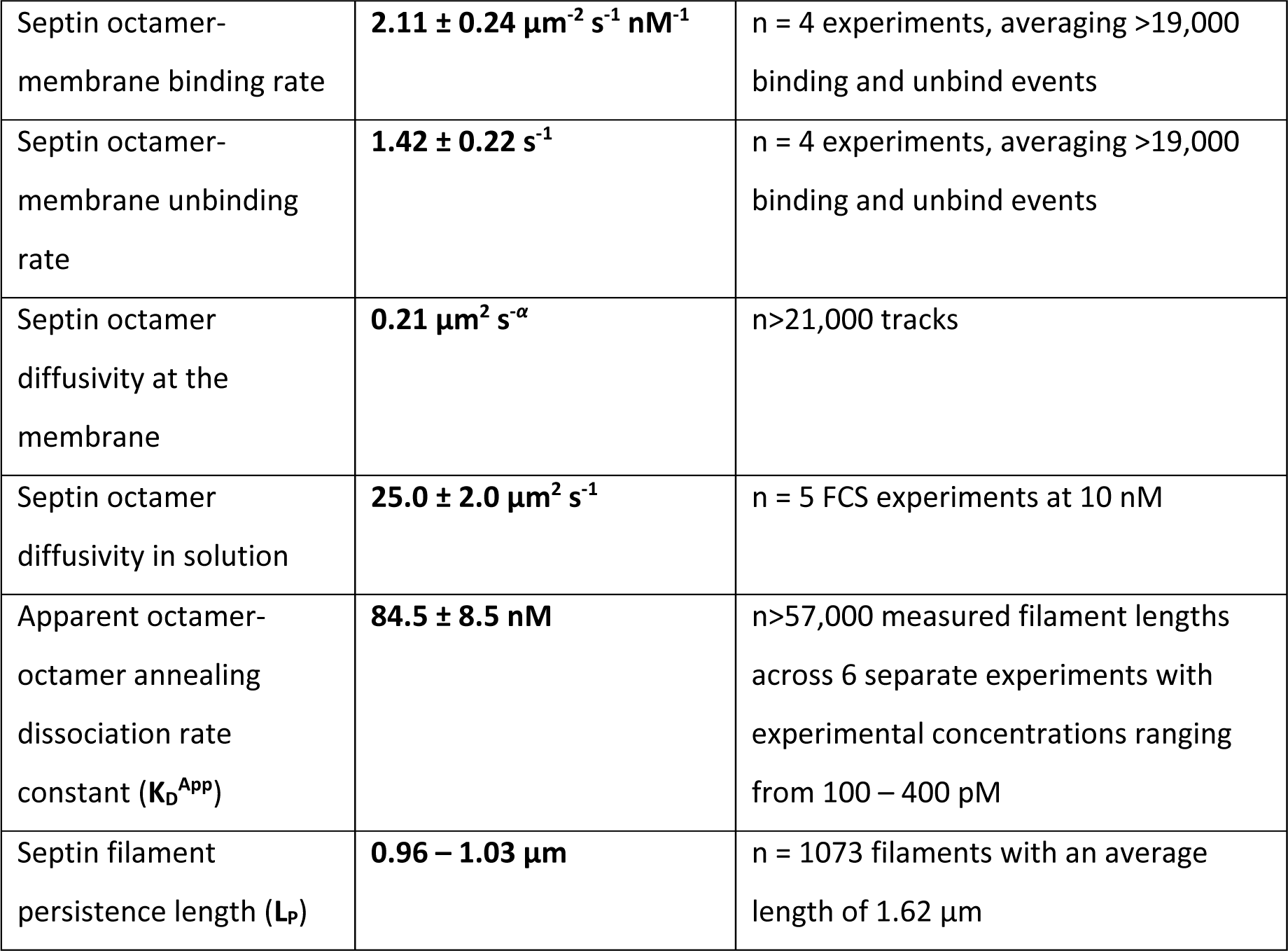
Measured reaction rates for septin polymerization.

We next assessed the steady state bulk adsorption of wild-type, polymerizable septins onto SLBs at a range of concentrations from 0.1 nM to 10 nM. Predictably, steady state adsorption levels increased with higher septin concentrations (**Fig 3B**). Interestingly, the fraction of total septins that adsorbed onto the membrane at steady state at all tested concentrations fell within a narrow range (**Fig 3C**). We note that at concentrations tested in our reconstitution assay, we did not detect pre-formed filaments in solution associating with the membrane. Likewise, we did not detect filaments dissociating from the membrane into solution. This is consistent with the observation that septin polymerization in fungal cells is localized exclusively to the plasma membrane (Bridges and Gladfelter, 2015; Bridges et al., 2014).

### Measuring septin octamer diffusivity on supported phospholipid bilayers

We next considered the mobility of septin octamers at the membrane and in solution. To measure the diffusivity of septin octamers on membranes, we incubated recombinant non-polymerizable, cdc11α6-capped octamers on phospholipid bilayers and tracked their displacement over time. The mean squared displacement (MSD) was determined by fitting a linear least squares regression in log-log space (**Fig 3D**). This fit corresponded to a septin diffusion constant of **0.21 μm^2^** *s*^−*α*^ (**Table 1**) and was slightly subdiffusive (α =0.86). Because freely diffusing octamers in solution are included in the simulations (see **Methods**), we determined diffusivity of septin octamers in solution by FCS. The diffusion constant in solution was **25.0 ± 2.0 μm^2^ s^-1^** (**Table 1**) similar to previously published results (Bridges et al., 2014).

### Septin octamers interact with nanomolar affinity

We next determined the binding affinity between septin octamers, which we term “annealing affinity” specifying the end-on interactions between the terminal subunits of the octamers. For an isodesmic polymer, the probability of fragmentation (dissociation) between any two septin octamers is equal irrespective of their position within the filament (**Fig 1A**). Therefore, the apparent dissociation equilibrium constant (**K_D_^app^**) between subunits of an isodesmic polymer can be determined as a function of the average polymer length (***L***) at steady state and the total concentration of subunits (Oosawa and Kasai, 1962; Romberg et al., 2001). We measured the membrane-bound septin concentration (i.e. density) by fluorescence intensity and this octamer density per area at the membrane is then multiplied by the height of septin octamers (4 nm), a previously described method to estimate the membrane-bound concentration in molarity (Bertin et al., 2008; Wu et al., 2011). The average filament length was determined from the fitted exponential to the length distribution (**Fig 2D**). At a range of septin concentrations we estimated the apparent dissociation equilibrium constant between octamers annealed end-to-end to be **84.5 ± 8.5 nM** (**Table 1**). This estimate is higher than what has been previously measured by in solution by FRET (25-30 nM) although still well within an order of magnitude (Booth et al., 2015). For probable reasons why these values differ, see the **Discussion**.

### Septin filaments have micron-scale flexibility

To determine the mechanical properties of septin filaments, we measured the persistence length (**L_P_**) of septin filaments on supported phospholipid bilayers by fitting an exponential to the extracted cosine correlation data of filament tangent angles (**Fig 3E**) (Graham et al., 2014). This semi-automated analysis determines the arc length above which the tangent angle becomes uncorrelated and has been used to measure the persistence length of actin and microtubule filaments (Gittes et al., 1993). The exponential fit to the cosine correlation corresponded to a persistence length of **0.96 – 1.03 μm** (**Table 1**).

### Brownian dynamics simulations of isodesmic polymerization

Measured biophysical properties were integrated as parameters in Brownian dynamics simulations based on a two-dimensional physical model of isodesmic polymerization (**Fig 2G, 4A**). In the model, octamers are represented as divalent spheres which can diffuse and anneal when binding sites are oriented, polymerizing into filaments over time (**Fig 4B**). Single octamers can bind to or unbind from the membrane but only participate in excluded volume and annealing interactions when they are membrane bound. Octamers “in solution” (gray in **Fig 4A**, not shown in **Fig 4B**) freely diffuse at a rate 100-fold faster, and do not interact with one another. An octamer incorporated into a filament does not unbind the membrane unless it dissociates from the filament. Thus, a key feature of the model is that membrane dissociation is exclusive to single octamers. Supporting these experimental observations, a theoretical analysis of cooperative assembly suggests that increasing the interface by a factor of two increases the equilibrium association constant by orders of magnitude (Erickson, 1989). Thus, even dimers of octamers should be essentially irreversibly bound to the SLB. The model only includes end-on annealing and excluded volume interactions among membrane-bound octamers and filaments, without consideration for filament pairing, crosslinking, or layering because we could not detect these events in reconstitution experiments with recombinant septins where we derived the parameters. During the course of a simulation, we measured the mean filament length and octamer-membrane adsorption, determined the filament length distribution, and assessed filament packing arrangements.

**Figure 4.**
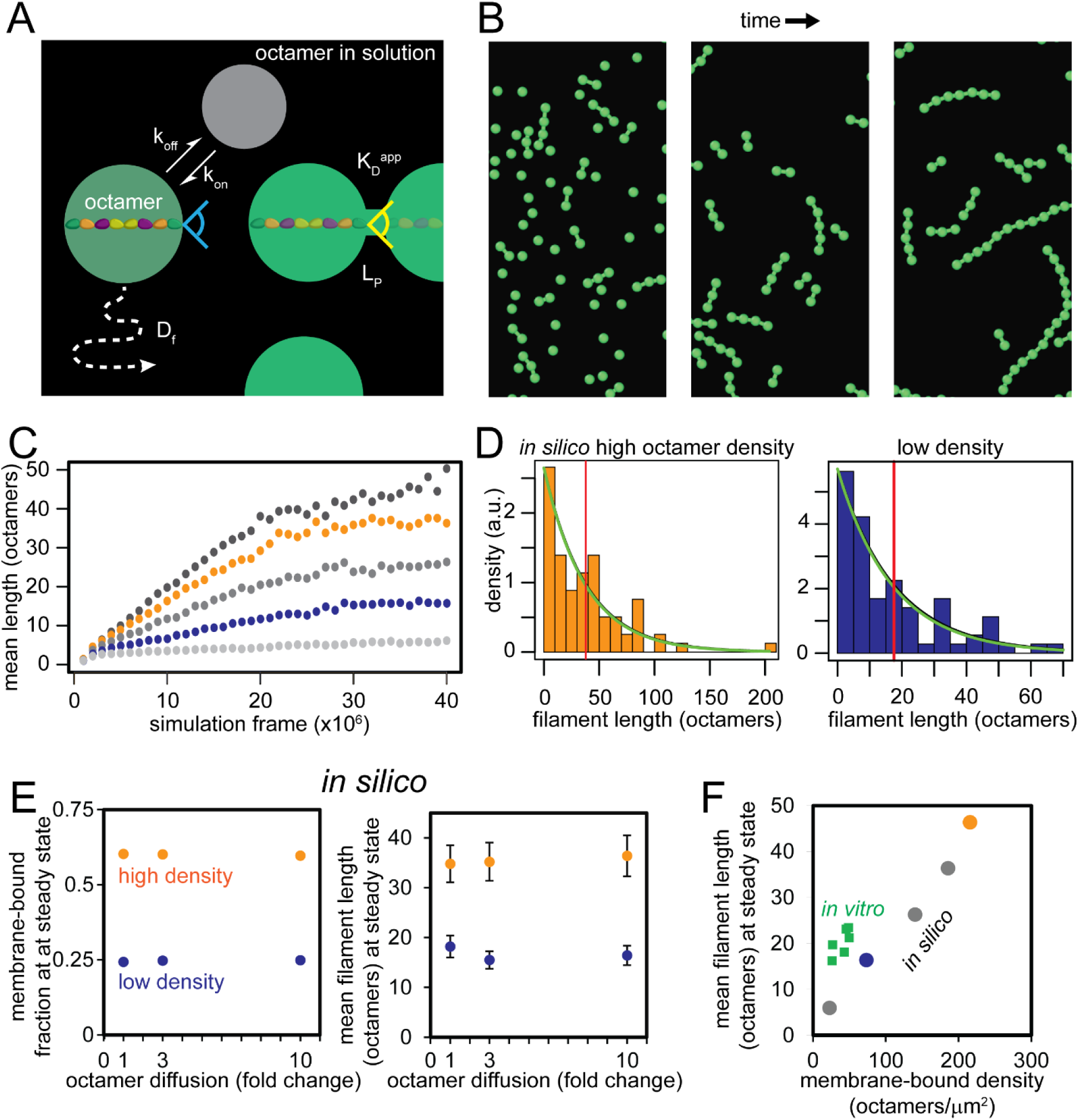
Parameterizing a molecular dynamics model of isodesmic polymerization. A. Octamers are represented as circles in the model. Octamers freely diffuse (**D_F_**), and can bind and unbind the membrane (k_on_ and k_off_ respectively). Octamer-octamer interactions occur at rate proportional to likelihood of octamers being aligned (see Methods). Octamer-octamer dissociation was scaled to reflect the apparent dissociation rate constant measured *in vitro* (**K_D_^app^**). The angle of potential octamer-octamer interactions was determined by the measured persistence length (**L_p_**). Octamers can only polymerize at the membrane, and while incorporated in polymers cannot unbind from the membrane. B. *In silico* septins polymerizing over time. Octamers “in solution” not depicted. C. Plotted mean filament lengths over the course of the simulation “seeded” with varying proportions of octamers starting on the membrane at the simulation. Low-density (in blue, fraction on membrane = 0.5) and high-density (in orange, fraction on membrane = 0.9) indicated for subsequent parameter sweeps (see **Fig 4D-F, 5F-G, 7D-F, 8B-C**). D. *In silico* filament length distributions at steady state for high (orange) and low (blue) densities. Green line indicates best exponential fit. Red line is the average *in silico* filament length determined from the fit. E. Plotted membrane-bound fraction and average filament length at steady state varying octamer diffusivity *in silico*. F. Plotting *in silico* mean filament length at steady state as a function of the steady state octamer density (calibrated by starting membrane-bound concentration, see **Fig 4C**). Also plotted are six representative examples of average *in vitro* filament lengths at steady state at indicated membrane-bound octamer densities (calibrated from single octamer fluorescence, see **Fig 1E**).

Reconstitution experiments *in vitro* typically take several hours to reach steady state conditions (**Fig 2C**). To mimic experimental conditions in the model, we ran simulations starting with no membrane-bound octamers. Over time octamers bound the membrane and began to polymerize, but simulations recapitulating *in vitro* conditions (starting with no membrane-bound octamers) were prohibitively slow due to computational limitations, never reaching “steady state” - which occurs when average filament length plateaus - in reasonable time frames. To mitigate the computational limitations, simulations were run with two alterations. First, simulations were “seeded” with membrane-bound octamers at their start. Predictably, this accelerates initial polymerization by removing the time lag to initiate polymerization, allowing simulations to reach steady state (**Fig 4C**). Seeding simulations with membrane-bound octamers also facilitates more precise control of the membrane-bound octamer density at steady state, important for comparing results from the model and *in vitro* experiments. At different membrane-bound octamer densities *in silico* polymer lengths were exponential distributed (**Fig 4D**) consistent with isodesmic polymerization on the membrane. Seeding the membrane also pushes the system into a regime that is not diffusion-limited, since octamers are close enough together that they can easily find each other during an average dwell time across a range of diffusion rates. Secondly, simulations could be accelerated by increasing octamer diffusion rates, which did not affect steady state properties such as membrane-adsorption and average filament length (**Fig 4E**).

Filament lengths measured *in silico* were directly proportional to the membrane-bound octamer density (**Fig 4F**). However, we found that *in silico* polymer lengths were slightly shorter compared to *in vitro* average lengths when normalized for octamer density (**Fig 4F**). We consider two possible reasons for this discrepancy. One possibility is that this reflects a finite size effect of the physical model. To reduce simulation times, the model is limited to 5000 total particles (octamers) diffusing in a continuous plane with periodic boundaries (e.g. particles that diffuse out one boundary will emerge from the opposite). Second, the *in silico* annealing angle between octamers may underrepresent reality. The angle in the model is a function of the filament flexibility (see **Methods**). A larger tolerated binding angle in the model would increase the probability of annealing interactions and increases filament lengths. Thus, the model provides a reasonable in silico approximation of the intrinsic septin assembly process. Our next goal is to use yeast cell extracts and the model to interpret how different septin regulators could impact these specific reactions to form varied higher-order septin assemblies seen in cells.

### Septins from cell extracts have a higher apparent annealing affinity and polymerize into rings

To determine how septin-regulatory proteins influence reactions, we analyzed septin filaments in whole cell extracts prepared from either wild-type or mutant *S. cerevisiae* cells on SLBs. Extracts were prepared by adapting previously published protocols from studies investigating F-actin and microtubule assembly (Bergman et al., 2018; Michelot and Drubin, 2014). Asynchronous cultures of wild-type haploid cells expressing GFP-Cdc3 from the endogenous locus were harvested during log-phase growth for extract preparation. Septins from cell extracts quickly polymerized (minutes) on SLBs into dense arrays of locally aligned filaments that appeared relatively rigid (**Fig 5A**). However, their density precluded segmentation necessary to quantify filament properties such as length and flexibility by TIRF microscopy. In contrast, septins from more diluted preparations polymerized into well-spaced filaments on the SLB and could be individually segmented and measured (**Fig 5A**). Septin adsorption increased with time reaching steady state after 3 hours (**Fig 5A & B**). At steady state, filament length distributions were exponential (**Fig 5C**), suggesting that septin polymerization from extracts is also isodesmic. Interestingly, filaments from cell extracts were twice the length of recombinant filaments at steady state (**Fig 5D**) after normalizing for the membrane-bound concentration as determined by single octamer fluorescence (**Fig 5E**). This could indicate octamers from extracts have a higher annealing affinity. Alternatively, unknown factors (including regulators) may promote filament extension (perhaps by expanding the angles of contact between octamers sufficient for annealing) or stabilize filaments from fragmentation.

**Figure 5.**
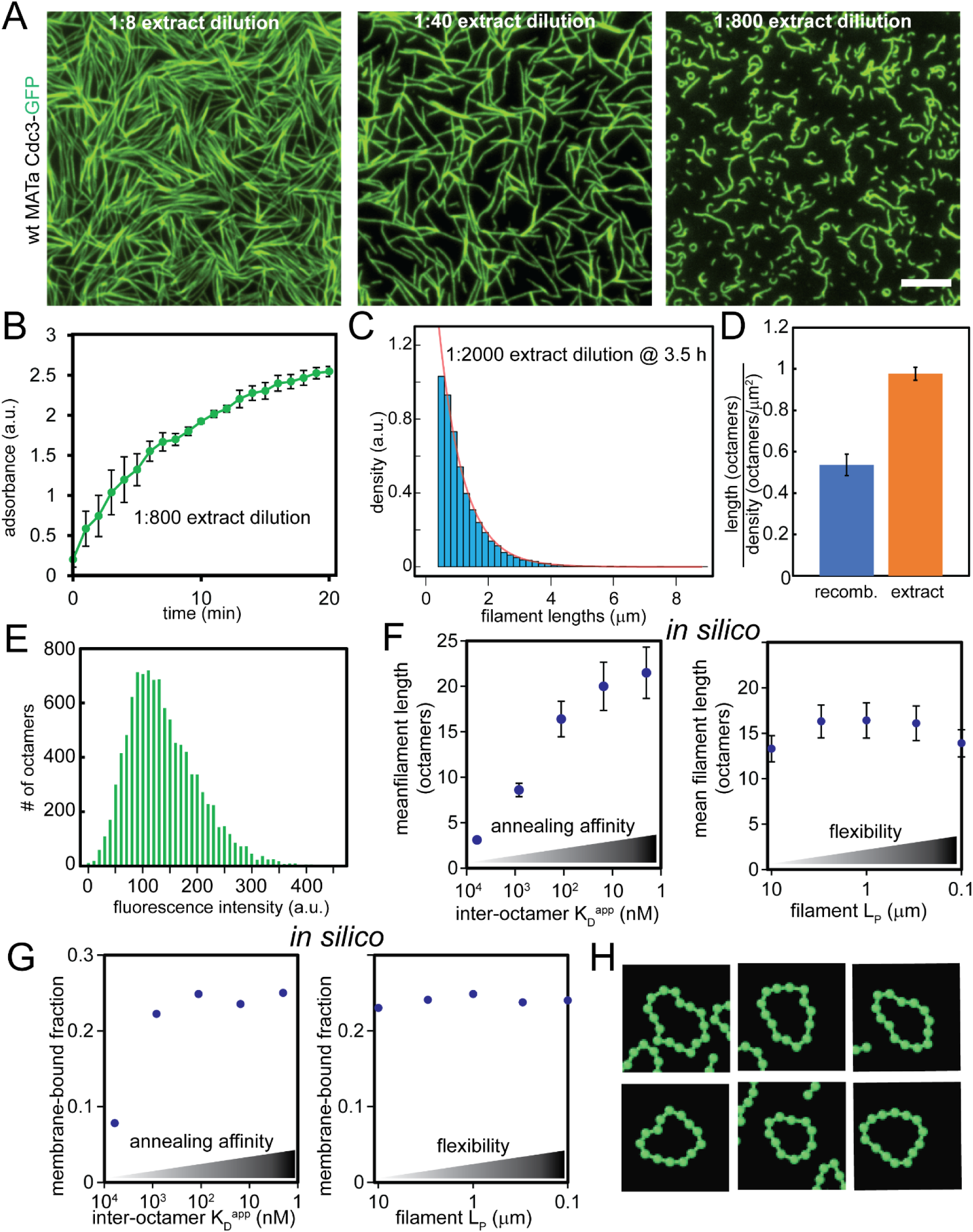
Septins from cell extracts undergo isodesmic polymerization. A. Extracts from wild-type MATa *S. cerevisiae* cells expressing Cdc3-GFP were incubated at indicated dilutions in reaction buffer (1.3 mg/ml BSA, 50 mM HEPES pH 7.4, 1 mM BME) on supported planar phospholipid bilayers (75% DOPC: 25% Soy PI). For dilutions higher than 1:8, extracts were first diluted into septin storage buffer (200 mM KCl, 50 mM HEPES pH 7.4, 1 mM BME) before a 1:4 dilution into reaction buffer. TIRF microscopy images acquired at steady state. Scale bar, 5 μm. B. Average fluorescence of septin filaments from wild-type budding yeast extracts over time (1:800 final cytoplasmic dilution, n = 3 separate extracts) assessed by fluorescence over background. Error bars represent standard error of the mean. C. Representative filament length distribution histogram of septin filaments at steady state with a fitted exponential (magenta line). D. Filament length (in octamers) of recombinant septins (>57,000 filaments from 6 experiments), and septins from extracts (>21,000 filaments from 3 experiments) relative to their density (octamers/μm^2^). Error bars represent standard error of the mean. E. Representative fluorescence intensity histogram of single septin octamers from cell extracts. Extracts were diluted 1:40,000 in septin storage buffer and incubated on plasma-cleaned glass. Non-mobile, individual fluorescent puncta adsorbed onto the glass were quantified. F. Plotted steady state mean filament length from simulations starting with a low membrane-bound fraction varying annealing affinity (left) and filament flexibility (right). After each simulation reached steady state, filament lengths fluctuated about a mean length. Error bars represent standard error of the mean once simulations reached steady state. G. Plotted steady state octamer-membrane adsorbance from simulations as in **F.** H. Representative ring-like filaments polymerize *in silico* when increasing both annealing affinity (K_D_^app^ = 2 nM) *and* filament flexibility (L_P_ = 100 nm).

One notable observation from diluted extracts was that filaments occasionally circularized into symmetrical rings approximately one micrometer in diameter (**Fig 5A**), reminiscent of rings at incipient buds and the split-rings at the yeast bud neck (Haarer and Pringle, 1987; Kim et al., 1991). Recombinant septin filaments never circularized into rings even at low densities, suggesting there are unique properties of filaments from extracts and/or the presence of regulators in extracts promote ring assembly. The differences between the two reconstituted assemblies (recombinant protein versus extract-based) were striking and indicate that factors in extracts substantially change intrinsic properties of septins.

### Model predicts ring-like filaments emerge from modulating combinations of parameters

We returned to the model to explore which parameters can be modulated to produce the distinct assembly features seen in extracts. We hypothesized that longer filaments from extracts may simply result from a higher octamer annealing affinity/decreased fragmentation rate, whereas ring formation could be modulated through filament flexibility, but how these features may interact is not intuitively obvious. Simulations were run at a low initial membrane-bound fraction to replicate low density polymerization conditions *in vitro*. We first adjusted *in silico* inter-octamer annealing affinity to assess how this parameter affects membrane adsorbance and polymer lengths at steady state. *In silico* filament lengths at steady state increased with a higher annealing affinity (**Fig 5F**), although adsorption changed little with altered annealing affinities in the model (**Fig 5G**). However, this effect was more subtle than we expected as the average *in silico* polymer length increased only 30% from the default annealing affinity in the model (equivalent to a **K_D_^app^** of ∼113 nM) to the highest annealing affinity we were able to test (**K_D_^app^** of ∼2 nM). Filaments polymerized from extracts *in vitro* were approximately twice as long as recombinant filaments (**Fig 5D**), suggesting a higher annealing affinity alone is unlikely to explain the differences in filament length between the recombinant and extract-based systems.

We next assessed how modifying filament flexibility (i.e. persistence length, **L_P_**) affected steady state properties. We predicted increased filament flexibility would promote filament circularization, thereby restricting filament extension. However, at low initial membrane bound fractions, altering flexibility alone did not noticeably affect mean length or adsorption (**Fig 5F & 5G**). Moreover, more flexible *in silico* filaments (equivalent to an **L_p_** of 0.1 μm) rarely circularized, contrary to our predictions. These results suggest that modulating flexibility alone is insufficient to promote ring formation.

We reasoned that regulators in extracts likely modulate multiple aspects of assembly simultaneously, and that a higher annealing affinity coupled with increased flexibility promotes assembly of rings. Indeed, increasing *both* the octamer annealing affinity and filament flexibility in the model promotes rings *in silico* (**Fig 5H**). Synthesizing the results from extracts and simulations suggest that distinct higher-order septin structures (such as rings) may result from regulators modulating multiple biophysical features (e.g. annealing and flexibility) of assembly either through direct binding and/or post-translational modifications.

### Reconstituted septin filaments polymerized from cell extracts are paired

Computational modeling indicated a much higher annealing affinity between octamers was insufficient to explain why filaments polymerized from extracts were twice as long as recombinant septin filaments (**Fig 5D**). What could explain the discrepancy in filament lengths between the two systems? A plausible explanation emerged when comparing fluorescence between the extract and recombinant filaments. The number of octamers by filament fluorescence versus the number expected by filament length are comparable for recombinant septins (**Fig 1G**). This suggests recombinant septin filaments were unpaired. Conversely, extract filaments, when normalized by single octamer fluorescence (**Fig 5E**), had twice as many octamers by fluorescence relative to filament length (**Fig 6A**), suggesting extract filaments are paired. Imaging septin filaments formed on supported phospholipid bilayers by scanning electron microscopy (SEM) confirmed that filaments polymerized from extracts were paired, whereas recombinant septin filaments were not (**Fig 6B**). Note that paired filaments were not exactly double the width of unpaired filaments due to the palladium-gold coating which has a thickness about the width of an unpaired filament (∼4 nm). It is unclear whether filament pairing occurs concurrently with polymerization or if one filament “templates” the other. We hypothesize pairing contributes to the long filaments in extracts because it reduces the probability of fragmentation since subunits within both conjoined polymers would have to dissociate in close proximity.

**Figure 6.**
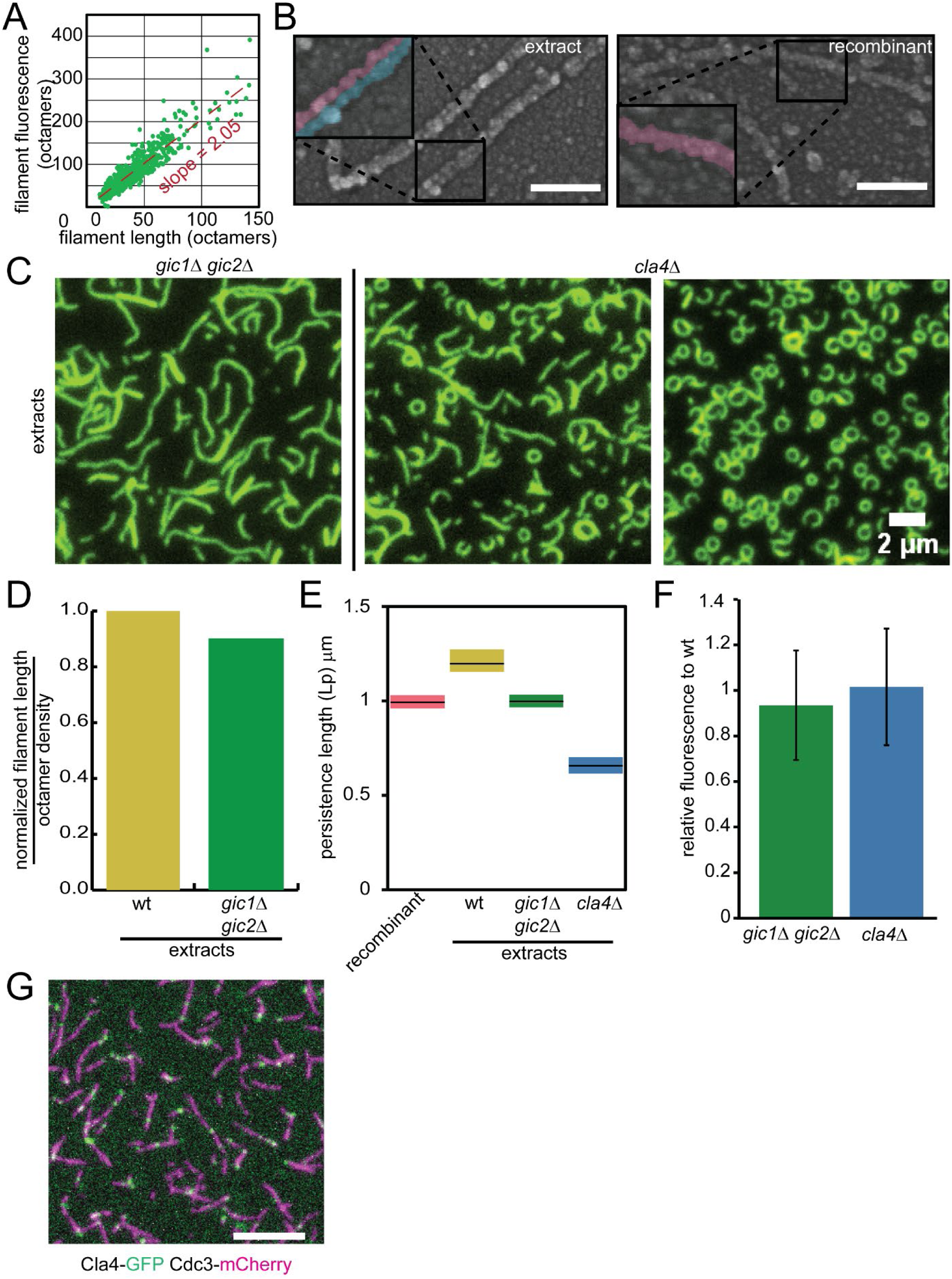
Analyses comparing filaments polymerized from mutant extracts. A. Scanning electron micrographs of septin filaments polymerized from recombinant protein (left) and wild-type cell extracts (right) on supported planar phospholipid bilayer. Inset panels display colorized traces of unpaired and paired filaments. B. Plot comparing length (in octamers) vs fluorescence (in octamers) of filaments polymerized from wild-type cell extracts. C. Filaments polymerized from asynchronous *gic1*Δ *gic2*Δ and *cla4*Δ mutant extracts at the same dilution on supported planar phospholipid bilayers and imaged by TIRF microscopy. Right *cla4Δ* extract panel is another representative field of filaments forming rings. Time point = 30 min. Scale bar, 2 μm. D. Comparing filament lengths as a function of octamer density between wild-type and *gic1Δ gic2Δ* mutant extracts. (Measurements from 3802 and 3855 filaments, respectively.) E. Persistence length (by cosine tangent angles as in Figure 3D) of recombinant septin filaments, and filaments polymerized from wild-type, *gic1*Δ *gic2*Δ and *cla4*Δ extracts. Colored bars denote the L_p_ range, whereas the black line is the mean. At least 1500 filaments from each genotype were measured with an average length > 1 μm. F. Normalized fluorescence of filaments (per unit length) polymerized from *gic1*Δ *gic2*Δ and *cla4*Δ mutant extracts (measurements of 1696 filaments and 2224 filaments, respectively) relative to filaments polymerized from wild-type extracts (measured 808 filaments). Error bars are standard error of the mean. Wild-type filament fluorescence is normalized to 1. G. Filaments polymerized from extracts harvested from wild-type haploid cells expressing Cdc3-mCherry and Cla4-GFP.

**Figure 7.**
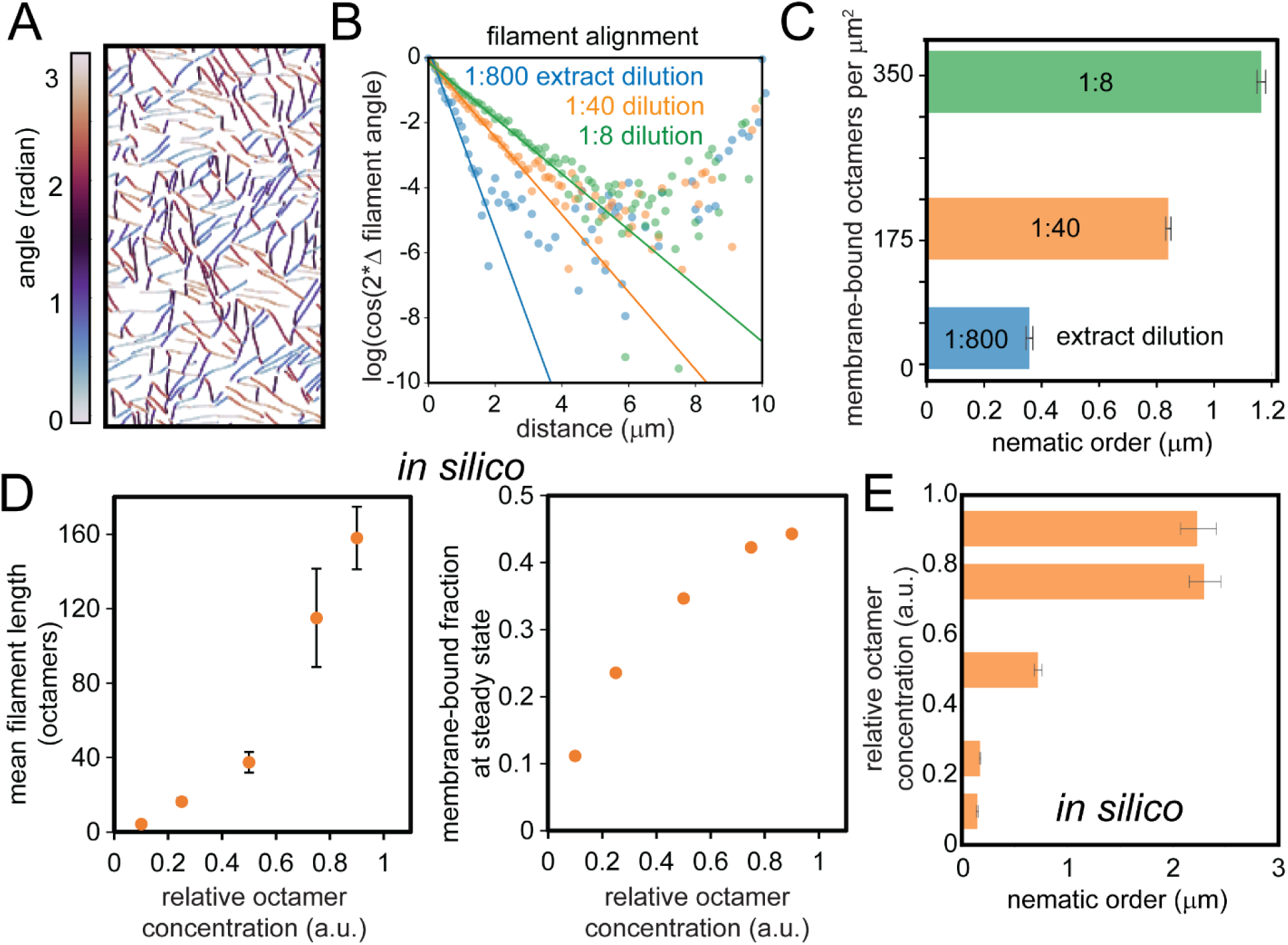
Filament alignment as function of membrane-bound septin concentration. A. Segmented filaments color coded based on relative angle (radians) from a 1:40 extract dilution (final) incubated on an SLB (as in **Figure 5A**). B. Plotted log-transformed average cosine correlation of angle differences as a function filament distance at varying incubated extract dilutions. Angle differences from 10^6^ randomized filament pairings per image (with an area >7700 μm^2^) times five representative images (5×10^6^ pairings total) at each extract titration. C. Nematic filament correlation as a function of membrane-bound octamer concentration at varying indicated incubated extract dilutions. Nematic order is extracted from the fit in **Figure 7B.** Error bars represent error in the cosine correlation fit. D. (Left) Plotting mean filament length from simulations starting as a function of total octamer concentration (membrane-bound + unbound). (Right) Plotting *in silico* octamer adsorbance as a function of total octamer concentration. E. Nematic correlation of *in silico* filaments as a function of total octamer concentration. Nematic order parameter analyses from 10^7^ *in silico* filament pairings at each tested density.

### Gic1, Gic2 and Cla4 tune septin filament flexibility but are not required for filament pairing

The fact that septin filaments from whole yeast cell extracts were twice as long as recombinant filaments, circularized into rings, and were paired shows cellular factors modulate distinct features of the septin polymer. Previous studies found that under certain conditions, recombinant septin filaments can be paired, possibly through coiled-coil interactions between the C-termini of Cdc3 and Cdc12 (Bertin et al., 2008). However, truncating the coiled-coil of Cdc3 or Cdc12 is lethal, precluding our ability to test this hypothesis in extracts (Finnigan et al., 2015; Woods et al., 2020). Alternatively, filament pairing could be mediated through septin regulators. Previous work with recombinant proteins demonstrated Gic1 can mediate septin pairing through its interactions with the inner-most septin subunit, Cdc10 (Sadian et al., 2013). Gic1 and its paralog Gic2 are functional homologs to the human Borg proteins (Joberty et al., 2001; Sheffield et al., 2003) and Cdc42 effectors that mediate cytoskeletal rearrangements during cell polarization, including septin recruitment to the cell polarity site (Brown et al., 1997; Daniels et al., 2018; Hall and Russell, 2004; Iwase et al., 2006). To assess their role in pairing in cells, we prepared extracts from mutants lacking Gic1 and Gic2.

We found filaments from *gic1*Δ *gic2*Δ double mutant extracts were similarly long as wild-type filaments (normalizing for membrane-bound concentration) at steady state (**Fig 6C, D**). Interestingly, filaments from *gic1*Δ *gic2*Δ extracts were more flexible than wild-type extract filaments (**Fig 6E**), suggesting Gic1 and Gic2 do tune filament flexibility. However, filaments polymerized from *gic1Δ gic2Δ* were still paired as assessed by filament fluorescence (**Fig 6F**), suggesting additional factors can mediate pairing in cells.

How else might septin filament pairing be regulated? Septins are post-translationally modified by phosphorylation, acetylation and even SUMOylation (Hernandez-Rodriguez and Momany, 2012). A leading candidate regulator is the p21-activated kinase (PAK), Cla4, which binds and phosphorylates septins (Versele and Thorner, 2004). In *cla4Δ* mutants, cells have wider bud necks with elongated buds and mis-localized septin structures (Cvrckova et al., 1995; Gladfelter et al., 2004; Longtine et al., 2000).

In reconstitution experiments with wild-type extracts expressing Cla4-GFP from the endogenous locus, Cla4 decorated septin filaments (**Fig 6G**). Strikingly, septins from *cla4Δ* mutant extracts were considerably more flexible than filaments from wild-type extracts (**Fig 6E**), and often formed fields of rings of uniform dimensions (**Fig 6C**), suggesting Cla4 inhibits ring formation possibly through its kinase activity. Nonetheless, filaments from *cla4Δ* extracts were still paired as assessed by filament fluorescence (**Fig 6F**). The abundant rings observed in *cla4Δ* extracts is consistent with our predictions from the model where increased flexibility coupled with a high annealing affinity (likely due to pairing in extracts limiting filament fragmentation) promotes filament circularization. In sum, Gic1 & Gic2 and Cla4 all contribute to filament rigidity, whereas only Cla4 impacts ring formation in extracts. However, neither Gic1 & Gic2 or Cla4 were necessary for filament pairing. It will be necessary for future work to disentangle the regulatory mechanisms governing filament pairing.

### Filament alignment in extracts is tuned by filament density

Extract-based experiments discussed thus far involved diluting extracts before their incubation with lipid bilayers to ensure filaments are well-spaced enough to analyze individual filament properties (e.g. flexibility and length). However, in cells, septin filaments are packed tightly at the polarity site and the bud neck, and such spatial constraints may influence the organization of filaments. Septins from more concentrated extracts incubated on supported lipid bilayers polymerized into densely packed networks of locally aligned filaments (**Fig 5A**) more reminiscent of aligned and clustered filament arrays observed at the bud neck (Ong et al., 2014). To quantify filament alignment as function of the membrane-bound septin concentration, we used a method that was originally developed to measure the organization of nematic liquid crystals (Copenhagen et al., 2020). First, we measured the relative filament angles (**Fig 7A**) at varying extract titrations on the supported lipid bilayer. We then plotted the average cosine of the filament angle differences as a function of the distance between measured points. The nematic order parameter (in distance units) is derived from the exponential fit (**Fig 7B**), and further explained in the **Methods**. The higher nematic order parameter value, the greater the degree of local alignment. The nematic order parameter analysis on experiments with wild-type whole cell extracts indicated that local filament alignment is correlated with the membrane-bound septin concentration (**Fig 7C**). Why are more densely packed septin filaments more ordered? One possibility is that filament polymerization in a crowded environment (such as the plasma membrane) promotes order by sterically constraining filaments into local zones of alignment. Alignment arising from being sterically constrained also suggests that filaments do not appreciably crisscross each other on SLBs, supporting the notion that septins are not “layering” in our assay. To test the crowding hypothesis, we analyzed *in silico* filament alignment from simulations run across a range of different octamer concentrations. Higher octamer concentrations increased lengths, adsorption, and filament alignment (**Fig 7D & E**), suggesting each of these steady state features are correlated in the model. In sum, results from the extracts and from the model suggest the degree of filament alignment can be in part be specified by the membrane-bound septin concentration changing through time. Additionally, the rate of accumulation at the membrane could influence the geometrical properties of the higher-order assembly, consistent with results observed on mica surfaces using atomic force microscopy (Jiao et al., 2020).

### Reducing filament flexibility promotes local filament alignment

We observed two additional higher-order assembly features in dense networks that may be interrelated and could impact alignment. Filaments in densely packed networks from extracts appeared less flexible compared to sparsely arranged filaments (**Fig 5A**). We were unable to estimate filament flexibility from TIRF microscopy images due to considerable filament overlap. However, measuring the flexibility of filaments in densely packed networks from SEM images corresponded to a persistence length of 6.88 ± 0.8 μm (65 filament segments, R = 0.999) (**Fig 8A**). In contrast, the persistence length of well-spaced extract filaments imaged by SEM was 1.63 ± 0.12 μm (23 filaments, R = 0.999), similar to **L_P_** measurements from TIRF microscopy (**Fig 6E**). It is unclear whether reduced flexibility observed in dense networks necessarily reflects inherent mechanical properties of the filaments (possibly through the function of concentrated regulators in the extracts) or is result of steric interactions between filaments in crowded environments which constrains filament directionality. Recombinant septins incubated on SLBs at high concentrations (>1 nM) polymerized into filaments that were not distinguishable from one another (**Fig 2B**) – unlike extract-based filaments – precluding our ability to test whether this held true in the absence of regulators. Instead, we returned to the model to investigate how filament flexibility impacts alignment in crowded environments, running simulations mimicking densely crowded conditions across a range of flexibility settings *in silico*. Consistent with the extracts, at a high filament density in the model, filament flexibility is anti-correlated with filament alignment (**Fig 8B**). Moreover, increased filament alignment in this dense regime promoted extension without impacting adsorption levels at steady state (**Fig 8C**). These results indicate that straighter filaments promote alignment and filament extension.

**Figure 8.**
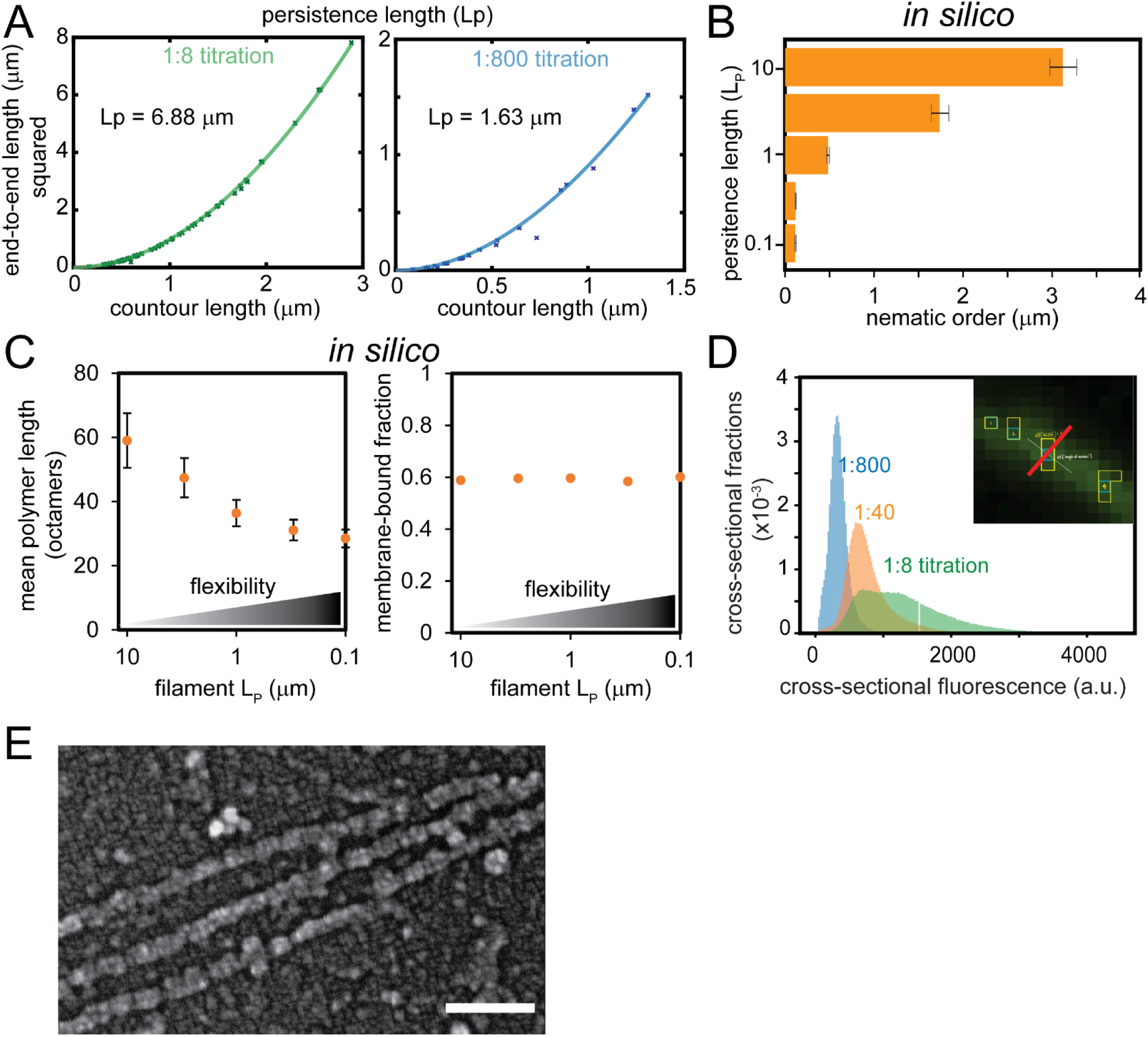
Reduced filament flexibility & filament bundling promote filament alignment. A. Persistence length measurements from SEM images with regressions fit to squared end-to-end lengths as a function of contour length. Plots compare analyses of filaments in densely packed networks (left; n = 65, average length = 1.01 ± 0.077 μm) and filaments in sparse arrangements (right; n = 23, average length = 0.504 ± 0.073 μm). B. Nematic correlation of *in silico* filaments at high-density as a function of filament flexibility (persistence length), measured from 10^7^ randomized pairings at each *in silico* flexibility setting. C. *In silico* filament length (left) and octamer-membrane adsorbance (right) at steady state as a function of filament flexibility in simulations starting with a high octamer-membrane density. D. Histogram of filament cross-sectional fluorescence at indicated extract dilutions. E. Scanning electron micrographs of closely aligned septin paired filaments polymerized from wild-type cell extracts on supported planar phospholipid bilayer.

The second observation from undiluted extract experiments is that filaments in dense networks appeared to be bundled. Although filaments overlapped in dense networks precluded our ability to completely segment individual filaments to quantify their overall fluorescence, the cross-sectional fluorescence of filaments was quantifiable, and indicated a sizeable fraction of filaments in dense networks were to two or three-times the cross-sectional fluorescence of filaments polymerized from diluted extracts (**Fig 8D**). SEM images of dense networks revealed that paired filaments do lie in close proximity with one another (**Fig 8E**). We conjecture bundling reduces capacity for filaments to flex in dense networks promoting their alignment. It will be necessary for future investigations to determine whether reduced filament flexibility in extracts is due to the concentration of regulators, filament bundling, or steric constraints during polymerization.

## Discussion

### Septin filament polymerization on planar lipid bilayers is isodesmic

Here we report that septin filament elongation is most consistent with isodesmic polymerization. This is based on three observations. First, septin filaments fragment (**Fig 1B & C**), which had been previously reported (Bridges et al., 2014; Khan et al., 2018). Spontaneous filament fragmentation is infrequent with cooperatively-formed polymers. Second, we could not find a threshold concentration below which filaments did not polymerize (**Fig 1D & H**), suggesting there is no critical concentration necessary for polymerization. This was coupled with the fact that septins polymerized on SLBs without a detectable lag (**Fig 1B & I**). Lastly, and most convincingly, septin filament length distributions are exponential at all testable concentrations (**Fig 1H, 2D & 2E, 5C**). In sum, these data suggest that septin filament polymerization is isodesmic on lipid bilayers and this is an intrinsic feature of the proteins.

### Septin filament fragmentation without ‘reptation’

Although isodesmic polymerization has seldom been described for cytoskeletal polymers, there is a documented instance in which actin polymerization is actually isodesmic. Actin in eukaryotic parasites, such as *Toxoplasma gondii*, is divergent from actin in other eukaryotes (Dobrowolski et al., 1997). Interestingly, there is no detectable polymerization kinetic lag in biochemical experiments using actin from *T. gondii,* which is atypical for actin (Skillman et al., 2013). Moreover, *T. gondii* actin filaments showed a wide distribution in lengths at steady state, atypical for a cooperative polymer. Thus, in some organisms, actin polymerization may be isodesmic.

It should be noted that an exponential distribution of filament lengths is not necessarily indicative of isodesmic polymerization. For example, when highly purified G-actin is incubated at very high concentrations sufficient to induce instantaneous polymerization, F-actin filament lengths are exponentially distributed (Sept et al., 1999). This stems from two causes. One is that at very high actin filament densities, filaments anneal end-to-end with one another, generating longer, interweaved filaments that can vary significantly in length. The second related cause is ‘reptation’ of polymers in a dense network where filaments slide along and collide with one another causing them to fragment at rate proportional to their length (Sept et al., 1999). Since reptation is not a factor where septin filaments are well dispersed on supported phospholipid bilayers, we infer that exponential distributions of septin filaments on planar lipid bilayers is due to isodesmic polymerization rather than filament ‘reptation’.

### Modeling isodesmic polymerization at the membrane

We constructed a Brownian dynamics model of septin polymerization that explicitly encoded octamers as divalent spheres that assembled into filaments via an isodesmic process; we specified that octamer-octamer binding occurred with a fixed energy, which leads to non-cooperative polymerization and filament fragmentation rates proportional to their lengths. The model incorporated experimentally determined values for the octamer-membrane affinity, octamer diffusion at the lipid bilayer, octamer-octamer annealing affinity, and filament flexibility. *In silico* filament lengths were exponentially distributed (**Fig 4D**), and bulk septin adsorption increased with membrane-bound octamer density (**Fig 4F**). These results are consistent with *in vitro* observations, supporting the notion that septin polymerization is isodesmic.

The simulations also allowed us to make mechanistic predictions about regimes of septin assembly inaccessible to experimental characterization, like very high-density, crowded filament fields. We observed that steric interactions lead to local alignment as a function of septin density at the membrane. We hypothesize that local alignment promotes the growth of longer filaments, which is consistent with our observation that a higher persistence length leads to longer filaments and enhanced local alignment (**Fig 8B**). All of the above predictions depend upon the inclusion of excluded-volume interactions in the model, which are computationally costly but biologically meaningful.

One measured parameter for the model, the apparent dissociation rate constant between octamers, differed in our estimates to one in the literature as analyzed by FRET (25-30nM) (Booth et al., 2015). Although this is a three-fold reduction in **K_D_^app^** reported here, these estimates are remarkably similar considering the numerous differences in which they were obtained. First, the fluorescent labeling strategy in Booth et al. required several point mutations throughout the recombinant septin octamer (2015). In Cdc11 these point mutations were C42F, C137A, C138A, with E294C becoming the exposed cysteine to control for labeling. These mutations do not block normal Cdc11 function in cells (de Val et al., 2013), but could affect octamer annealing affinity. Second, recombinant octamers in our assay were capped with Cdc11 fused at its C-terminus to a SNAP tag for fluorescent labeling. The C-terminal tag could also impact octamer-octamer interactions. Third, our estimation comes from experiments on supported planar bilayers whereas the estimates in Booth et al. were from interactions in solution (2015). Octamer interactions with the lipid bilayer could influence octamer-octamer interactions, perhaps exposing or masking different residues that are left exposed in solution.

In summary, we were able to use the model to predict how different parameters controlling septins may be modulated for septin regulation. It enabled us to examine how different parameters can interact to produce particular features of septin assemblies such as rings. As additional features are added to the model, including filament-filament interactions beyond steric constraints (e.g. explicit pairing or bundling), it will become an increasingly powerful tool for interpreting how different filament properties and arrangements result from tuning specific biophysical properties during assembly.

### Extracts from budding yeast as a tool to investigate regulation of filament polymerization

Cell extracts have been valuable tool for investigating cytoskeletal assembly. Reconstitution experiments using *Xenopus* egg extracts provided a foundational framework for studying microtubule spindle assembly (Dunphy and Newport, 1988; Lohka and Maller, 1985; Miake-Lye and Kirschner, 1985; Newport and Spann, 1987) and yielded key insights into actin assembly dynamics (Theriot et al., 1994). However, a limiting factor for most extract-based reconstitution systems including *Xenopus* is the lack of precision in controlling the extract contents genetically. Extracts from *S. cerevisiae* provide many the advantages of extract-based experimentation, but with the added benefit of leveraging the power of yeast genetics. Budding yeast extracts have been useful for studying microtubule and actin polymerization (Bergman et al., 2018; Michelot and Drubin, 2014). In this study, we employed a similar extract protocol to investigate the biophysical properties governing septin filament polymerization in budding yeast, where septins were first discovered and their biology is arguably best understood.

Reconstitution experiments utilizing wild-type cell extracts support the isodesmic model of septin filament polymerization (**Fig 5A,C**). Interestingly, core characteristics of isodesmic polymerization were unchanged despite filament pairing (**Fig 6A & B**). Septin filaments polymerized from extracts were twice as long per octamer density on supported lipid bilayers than filaments polymerized from recombinant protein (**Fig 5D**) suggesting filament pairing reduces (but does not block) fragmentation. Purified septins from yeast also polymerized into paired filaments (Frazier et al., 1998), indicating either a post-translational modification and/or stably bound regulator promote pairing. What else might promote filament pairing? It had been previously reported that Gic1 and Cdc42 mediate filament pairing *in vitro* (Sadian et al., 2013). However, filaments from *gic1Δ gic2Δ* extracts were still paired (**Fig 6F**) suggesting there are parallel pathway(s) mediating filament pairing.

Recent work investigating the architecture and arrangements of septin filaments in budding yeast via platinum replica EM have shed light on the intricate organizational transition of septin filaments through the cell cycle (Chen et al., 2020; Marquardt et al., 2020). After bud emergence, septin filaments aligned along the mother-bud axis are paired (Ong et al., 2014). Marquardt et al. demonstrate that the organization and pairing of axial filaments is dependent on the kinase, Elm1 (2020). These axial filaments are organized into a “zonal architecture” by single, unpaired septin filaments that run longitudinally around the circumference of the bud neck (Chen et al., 2020). Unpaired circumferential filaments are critically dependent on the regulators Bud3, a reported RhoGEF, and Bud4, an anillin-like protein. Later in the cell cycle just prior to cytokinesis, septins dramatically rearrange into paired, circumferential filaments sandwiching the actomyosin ring in the middle (Marquardt et al., 2020; Ong et al., 2014). Future extract experiments will enable exploration into how these different regulators combinatorially modulate filament pairing. We also anticipate the extract-based reconstitution assay will be a valuable tool to investigate how the cell-cycle impacts filament polymerization using well characterized *cdc* mutants (Hartwell et al., 1970; Hartwell et al., 1973).

### Filament flexibility and local alignment in densely packed networks

Septins incubated at low dilutions on supported lipid bilayers polymerized into highly flexible filaments with persistence lengths at about 1 μm (**Fig 3E**). Previously, it had been reported that septin filaments were more rigid, with a persistence length approaching 10 μm (Bridges et al., 2014). We suspect this discrepancy is due to the number of filaments measured previously (n = 10), whereas the analysis reported here was based on measurements of many more filaments (n > 1000) using a semi-automated analysis (Graham et al., 2014). The updated persistence length measurement is close to the optimal membrane curvature preference of septin filaments (Bridges et al., 2016; Cannon et al., 2019), underscoring how the mechanical properties of septin filaments may be well adapted to their function in distinguishing micron-scale membrane curvature. Septin filament flexibility can be modulated by the presence of particular regulators, such as Gic1 and Gic2 and Cla4 (**Fig 6E**). How is septin filament flexibility modulated at the subunit level? One possibility is that filament flexibility is tuned at the level of the octamer itself. Cla4 stably associates with Cdc10 and Cdc3, subunits within the core of the septin octamer (Versele and Thorner, 2004), and Cla4 decorated filaments during polymerization (**Fig 6G**). Cla4 may stiffen octamers directly by its association with core octamer subunits, or by its kinase activity.

An alternative possibility is that the interaction angle between octamers dictates filament flexibility. Septins are post-translationally modified by SUMOylation, acetylation, and phosphorylation (Hernandez-Rodriguez and Momany, 2012). Septin octamers can be capped with the non-essential septin subunit Shs1 in place of Cdc11, and Shs1 has multiple residues that are phosphorylated (Egelhofer et al., 2008; Meseroll et al., 2012; Meseroll et al., 2013). Previous studies *in vitro* found that incorporation of wild-type Shs1-containing octamers promotes filament circularization (Garcia et al., 2011). Intriguingly, phosphomimetic mutants of Shs1 at the NC-interface inhibited ring formation, and promoted filaments to be arranged in lattice-like networks (Garcia et al., 2011). An attractive hypothesis is that Shs1 phosphorylation by Cla4 inhibits ring assembly, possibly by biasing Cdc11-capped octamer incorporation into filaments. This could explain why septins from *cla4*Δ mutant extracts had an increased tendency to polymerize into rings (**Fig 6C**).

## Conclusion

Here we report that septin filament polymerization is isodesmic. Gleaning biophysical properties of septin filament polymerization from reconstitution assays, we parameterized a physical model of septin filament assembly at the membrane. Using whole cell extracts, we began to probe how different proposed septin regulators modulate these biophysical properties in the assembly of higher-order septin structures which we could recapitulate in the model. This work provides a quantitative foundation for understanding the intrinsic and extrinsic basis for the formation of septin higher-order assemblies.

## Materials and Methods

### Lipid mix preparation

Lipids (Avanti Polar Lipids) were mixed in chloroform solvent at a ratio of 75 mole percent dioleoylphosphatidylcholine (DOPC) and 25 mole percent PI (liver, bovine) sodium salt in a glass cuvette. For lipid mix preparations to incubated on silica microspheres, trace amounts (>0.1%) of phosphatidylethanolamine-N-(lissamine rhodamine B sulfonyl) (RH-PE) were added. Lipid mixtures were then dried with argon gas to create a lipid film and stored under negative vacuum pressure overnight to evaporate trace chloroform. Lipids were rehydrated in Supported Lipid Bilayer buffer (300 mM KCl, 20 mM Hepes pH 7.4, 1 mM MgCl_2_) at 37°C to a final lipid concentration of 5 mM. Lipids were then resuspended over the course of 30 minutes at 37°C, with vortexing for 10 seconds every five minutes. Fully resuspended lipids were then bath sonicated at 2-minute intervals until solution clarified to yield small unilamellar vesicles (SUVs).

### Phospholipid mix preparation

Lipid small unilamellar vesicles (SUVs) were prepared as previously described (Cannon et al., 2019). Lipids (Avanti Polar Lipids) were mixed in chloroform solvent at 75 mole percent dioleoylphosphatidylcholine (DOPC) and 25 mole percent PI (Soy plant) sodium salt in a glass cuvette. To test lipid bilayers for lipid diffusivity by fluorescence recovery after photobleaching, a trace amount (>0.05%) of phosphatidylethanolamine-N-(lissamine rhodamine B sulfonyl) (RH-PE) was added. Phospholipid mixtures were dried under argon gas to evaporate the chloroform leaving a lipid film, and then stored under negative vacuum pressure overnight. Lipids were rehydrated in aqueous Supported Lipid Bilayer buffer (300 mM KCl, 20 mM Hepes pH 7.4, 1 mM MgCl_2_) at 37°C to a final lipid concentration of 5 mM. Lipids were then vortexed for 10 s every five minutes over the course of 30 minutes at 37°C until fully resuspended. Resuspended lipids were bath sonicated at 2-minute on/off intervals until suspension clarified yielding lipid SUVs.

### Preparation of planar supported phospholipid bilayers

No 1.5H cover glass (Matsunami Cover Glass Ind., Inc) were cleaned with oxygen plasma for 15 min at maximum power (PE25-JW, Plasma Etch). Chambers modified from PCR tubes (Bridges et al., 2014) were adhered onto cleaned cover glass under UV light for 5 min (Norland optical adhesive, Thor Labs). SUVs were incubated at 37°C on cover glass within chambers, diluted in Supported Lipid Bilayer buffer (with additional 1 mM CaCl_2_) to a final lipid concentration of 1 mM μl for 20 minutes. Ultra-pure, distilled water (Millipore Sigma) was necessary for any solution used in preparation or experiments with planar supported lipid bilayers to obtain reproducible results. After vesicle fusion, supported lipid bilayers were washed six times with 150 μl of Supported Lipid Bilayer buffer (without CaCl_2_) to rinse away excess SUVs. Immediately prior to adding septins or extracts, lipid bilayers were rinsed six times with a 150 μl of Reaction Buffer (50 mM Hepes pH7.4, 1 mM BME, and 0.13% low fatty-acid BSA [product no. A4612, lot no. SLBM2472V, Millipore Sigma]). The product number and lot number of low fatty-acid BSA were important for reconstitution assay reproducibility.

### Protein purification

Recombinant *S. cerevisiae* septin protein complexes were expressed from a duet-cassette expression system in BL21 (DE3) *E. coli* cells and purified as previously described with minor modifications (see **Table 2** for plasmids used in this assay) (Bridges et al., 2016; Cannon et al., 2019). Transformed *E. coli* cells were selected on LB medium agar (2%) plates with ampicillin and chloramphenicol. Cells were cultured in 1 L liquid LB medium to an OD_600nm_ of 0.6-0.7 before addition of 1 mM isopropyl-β-D-1-thiolgalactopyranoside (ITPG; Thermo Fisher Scientific) to induce expression. Induced cultures were grown with gentle shaking for 24 h at 22°C before harvesting. Cultures were pelleted at 7500x relative centrifugal force (RCF/g) for 20 min at 4°C. Pelleted cells were then resuspended in lysis buffer (1 M KCl, 50 mM Hepes pH 7.4, 1 mM MgCl_2_, 10% glycerol, 1% Tween-20, 20 mM imidazole, 1 mg/ml lysozyme, and 1x protease tablet [Roche]) for thirty minutes on ice with 5 min intervals of 10 s vortexing. Lysates were kept chilled, and sonicated twice for 10 s with a one minute interval. Lysates were centrifuged in an SS-34 rotor at 20,000x RCF for 30 minutes at 4°C. Clarified lysates were passed over a Acrodisc PF syringe 0.8/0.2 μm filter (VWR) prior to incubation with HisPur^TM^ cobalt resin (Thermo Fisher Scientific; 2 ml per liter of *E. coli* culture) for one hour at 4°C with gentle rocking. Protein bound to the resin was washed four times with five resin-column volumes of wash buffer (1 M KCl, 50 mM Hepes pH 7.4, and 20 mM imidazole), then once with pre-elution buffer (300 mM KCl, 50 mM Hepes pH 7.4). Protein was then eluted off the cobalt resin with five resin-column volumes of elution buffer (300 mM KCl, 50 mM Hepes, 500 mM imidazole). Eluted protein was concentrated (if necessary) using 100 kDa cutoff Amicon Ultra centrifugal filters (EMD Millipore). Purified protein was dialyzed into storage buffer (300 mM KCl, 50 mM Hepes pH 7.4, 1 mM *β-*mercaptoethanol [BME]) in two steps using Slide-A-Lyzer G2 (20 kDa cutoff; Thermo Fisher Scientific), with 60 μg of Tobacco etch virus protease (Sigma-Aldrich) added in the dialysis cassette to cleave the histidine tag, for 24 h at 4°C. Protein was then run over 1 mL cobalt resin to remove high cobalt-affinity contaminants, the cleaved histidine tag, and the protease. Protein purity was assessed by SDS-PAGE followed by silver staining, and protein concentration was measured by a Bradford assay.

**Table 2.** Plasmids used in this study.

| Plasmid | Construct | Selection | Source |
| --- | --- | --- | --- |
| AGB501 | pMVB133 *S.c.CDC3/S.c.CDC11*-SNAP | cam | (Bridges et al., 2016) |
| AGB710 | pMVB128 6xHIS-TEV-*S.c.CDC12/S.c.CDC10* | amp | (Bridges et al., 2016) |
| AGB744 | pMVB133 *S.c.CDC3/S.c.cdc11α6<sup>E289R,Y291A,R292A</sup>*-SNAP | cam | (Bridges et al., 2016) |

### Yeast strain construction and culturing

Standard molecular genetic techniques and procedures were used to generate and passage yeast strains in this study. All strains used in this study can be referenced in **Table 3**. We received a strain expressing a Cdc3 construct with an internal GFP fusion from the endogenous *CDC3* locus from Daniel Lew (Duke University), described previously (Chen et al., 2011; Tong et al., 2007). We also received strains deleted for *CLA4* and *GIC1* plus *GIC2* from Daniel Lew (Duke University), which were previously described (Daniels et al., 2018; Kozubowski et al., 2008). To grow yeast cultures for whole cell extract preparation, ∼10^9^ cells from bench stocks grown to stationary phase were inoculated into 1 L of YEP + 2% dextrose medium. Cultures were grown at 24°C for 16-18h with shaking to mid log phase 0.5-1×10^8^ cells/ml before harvesting.

**Table 3.** Yeast strains used in this study.

| Strain | Relevant Genotype | Source |
| --- | --- | --- |
| AGY581 | MATa GFP-*CDC3:URA3* (DLY10055) | Daniel Lew |
| AGY636 | MATa mCherry-*CDC3:LEU2 CLA4*-GFP:HIS3 | This study |
| AGY696 | MATa GFP-*CDC3:URA3 gic1::KAN<sup>R</sup> gic2::HIS3* | This study |
| AGY698 | MATa GFP-*CDC3:URA3 cla4::NAT<sup>R</sup>* | This study |

All strains used in this study were in the YEF473 genetic background (*his3-Δ200 leu2-Δ1 lys2-801 trp1-Δ63 ura3-52*) (Bi and Pringle, 1996).

### Yeast cell extracts

Preparation of yeast cell extracts for investigating septin filament polymerization was adapted from a previously published extract preparation method for studying actin (Michelot and Drubin, 2014). Briefly, harvested cultures were pelleted at 7500xRCF for 15 min at 4°C. Pellets were rinsed 10 ml with ultra-pure, distilled water. Cells were rinsed again in 5 ml of water and transferred to a 10 ml capped syringe placed in a 50 ml conical tube for centrifugation. Pellets were then dispensed from the syringe directly into a mortar filled with liquid N_2_, and repeatedly crushed to a fine powder with a mortar while continuously bathed in liquid N_2_. Yeast cell powder was weighed, then kept on ice in a beaker. Following this, 100 μl of cold Extract buffer (100 mM Hepes pH 7.4, 10 mM BME) and 4 μl of protease inhibitor (Protease Inhibitor Cocktail Set IV, EMD Millipore) was added to every gram of cell powder. Powder, buffer, and protease inhibitor were mixed with a spatula and kept on ice until the powder thawed to a liquid extract. Extracts were then clarified by ultra-centrifugation in a TLA100.3 rotor at 264,360x RCF (70000 RPM) for 20 min. Supernatant was collected (avoiding the lipid layer at the top and pellet layer at the bottom) and ultra-centrifugated again for 10 min. Resulting extract supernatant were kept on ice, and could be used for up to 4 hours. Unused extracts were frozen in liquid N_2_ for later use. See **Table 3** for strains used in this study.

### Reconstituting septin filament assembly on planar supported phospholipid bilayers

Recombinant septins were diluted to a pre-reaction concentration in Septin Storage buffer (200 mM KCl, 50 mM Hepes pH 7.4, 1 mM BME, 0.13% low fatty-acid BSA), and then incubated on supported lipid bilayers, titrated to the final indicated, septin-reaction concentration in solution with 50 mM KCl, 50 mM Hepes pH 7.4, 1 mM BME, and 0.1% BSA. Extracts were incubated on supported lipid bilayers much the same way, with any necessary titrations in Septin Storage buffer.

### Fluorescence Microscopy

Total internal reflection fluorescence (TIRF) microscopy images were acquired using a Nikon TiE TIRF system equipped with a solid-state laser system (15 mW, Nikon LUn-4), a Nikon Ti-82 stage, a 100x Plan Apo 1.49 NA oil lens, and Prime 95B CMOS camera (Photometrics). Exposure times for all TIRF microscopy images was 80 ms.

Fluorescence correlation spectroscopy (FCS) of soluble septins incubated on supported lipid bilayers was performed on an LSM 880 laser scanning confocal microscope (Zeiss) equipped with Airyscan, multi-use argon laser (Lasos) and a C-Apochromat 40x/1.2 NA water immersion objective. FCS data, correlations and protein diffusion rates were analyzed using ZEN 2011 software (Zeiss).

### Image analysis

Analysis and preparation of microscopy images were processed in FIJI/ImageJ unless otherwise noted. Recombinant septin octamer fluorescence was assessed by TIRF microscopy after incubating recombinant septin octamers (between 20 and 200 pM final concentration) harboring nonpolymerizable mutant cdc11α6 (fused to a SNAP-tag and conjugated to Alexafluor488 dye) directly on plasma cleaned cover glass or on supported lipid bilayers. Puncta were thresholded based on their fluorescence above background with a dilated region of interest (ROI) to include any scattered light in quantification. Fluorescent puncta intensity above background was plotted in a histogram to determine the mode fluorescent intensity, which was used to calibrate fluorescent intensity of single septin octamers (containing *two* fluorophores). Septin octamer fluorescence from extracts was determined the same way, but only on cleaned cover glass, at a final dilution of 1:20000. Octamer fluorescence was calibrated regularly but did not substantially vary day-to-day.

To determine the septin-membrane on/off rates and diffusivity on planar supported phosopholipid bilayers, recombinant cdc11α6-SNAP octamers were incubated on lipid bilayers for at least one hour and then imaged by TIRF microscopy. Fluorescent puncta were tracked using the TrackMate plugin (v5.2.0) for FIJI. Puncta were detected with sub-pixel localization and median filter applied. The Simple LAP tracker algorithm provided X,Y,t coordinates for each track, which were used to determine octamer mean squared displacement (M.S.D.).

#### Determining M.S.D

Three separate experiments contained 35,532, 22,837, and 13106 tracked paths, respectively. Each path contained x,y values in microns with 0.08 second intervals. Non-mobile puncta and paths with less than three x,y values were excluded. The M.S.D. was calculated as:

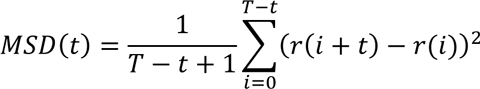

where t is the lag time, T is the total time of the path, and r(i) is the position of the particle at time i. The M.S.D. at the first 20 lags were calculated for each path, and the average and standard deviation of the M.S.D. was reported at each lag for each of the three experiments. Finally, the M.S.D. from the three experiments were averaged with the standard deviation for each propagated as:

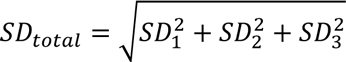

where *SD_i_* is the standard deviation of experiment i. To plot the average M.S.D. from the three experiments and the associated error, the lag times and average M.S.D. were log transformed using the natural log. To correctly represent the error in log-log space, the error on either side of the mean was calculated as:

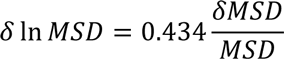

as reported previously(Baird, 1994). The gray shaded area in the plot (**Fig 3D**) shows the mean M.S.D. +− *δ* ln *MSD*.

#### Extracting the apparent dissociation equilibrium rate constant to approximate inter-octamer annealing affinity

Filament lengths were measured using the Ridge Detection plugin for FIJI. Compiled filament lengths were then plotted into a histogram using R. An exponential data fit to the lengths was manually parameterized by a floating lambda (λ) value reciprocal to the average filament length. Since filament lengths above the diffraction limit are exponentially distributed, we can extract the average length from an exponential fit. This is due to the memoryless property of exponential distributions – if the underlying distribution is exponential, any left-truncated (i.e. missing diffraction limited population) form will also be exponential, with the same rate parameter, λ. Therefore, the average length is determined to be the reciprocal of the rate parameter of the fit exponential distribution, 1/λ.

For an isodesmic polymer, the affinity can be determined by the average filament length at equilibrium and the total subunit protein concentration (i.e. septin octamer concentration) (Oosawa and Kasai, 1962). The apparent inter-octamer annealing dissociation rate constant (***K_D_^app^***) can be modeled with the following equation:

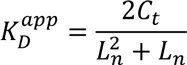

where ***C_t_*** is the measured octamer concentration and ***L_n_*** is the average length (Romberg et al., 2001).

The total septin concentration in a reconstitution assay does not reflect the “available” septin concentration (***C_t_***) for polymerization at the membrane. Since filament polymerization occurs predominantly at the membrane we only considered the membrane-bound septin concentration. The fluorescence intensity of single octamers was used to calibrate the septin octamer density at the membrane, which was averaged over multiple fields and multiplied to the entire area of the bilayer (2700 μm^2^). We were then able to transform the membrane-bound septin octamer density into a quasi-3D concentration assuming a septin octamer height of 4 nm (Wu, Honig, Nature 2011; Bertin et al., 2008).

We assessed the accuracy of our fluorescence-based concentration measurements by two different methods. First, we determined whether filament fluorescence correlated with filament length. We plotted filament length verse the filament fluorescence. Normalized for the fluorescence and length of single octamers (32 nm), we found that filament fluorescence correlated closely with length assuming recombinant filaments were unpaired and extract filaments were paired. This indicated that fluorescence measurements of single octamers by TIRF microscopy is a relatively accurate method by which to calibrate septin concentration by bulk fluorescence. We also measured the soluble septin octamer concentration in our *in vitro* reconstitution assays by FCS. The soluble fraction measured by FCS should equal the total septin concentration minus the membrane-bound septin fraction, and we found that the soluble concentrations were comparable to subtracted estimates by TIRF microscopy. These data suggest membrane-bound septin concentration measurements by TIRF microscopy are reasonably accurate.

#### Persistence length measurements

To determine the persistence length of reconstituted filaments imaged by TIRF microscopy, the cosine correlation of average tangent angles along the length of filaments were measured from single time points, and fitted to an exponential to determine the persistence length using a MATLAB GUI previously published (Graham et al., 2014). Cosine correlations were unreliable for measuring the persistence length of filaments imaged by scanning electron microscopy (SEM) due to the scale of the images, and generated poor exponential fits to the data. Instead, we determined persistence length of filaments imaged by SEM using a regression of filaments’ end-to-end distance as function of their contour length, which is an additional output of the same MATLAB GUI (Graham et al., 2014).

### Nematic order parameter

To measure local regions of alignment, we first quantify the orientation angle field of each image using the ImageJ Ridge Detection plugin. Using this orientation angle field, we calculate the spatial correlation function of the nematic orientation angle (Li et al., 2019):

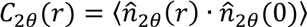

where 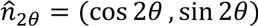. We use 5×10^6^ pairs of pixels (within filaments) in each image to calculate *C*_2*θ*_(*r*). We then fit the following function:

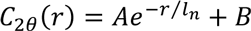

to obtain the nematic correlation length, *l*_*n*_.

### Scanning electron microscopy

For images acquired by scanning electron microscopy (SEM), filaments polymerized from recombinant protein or extracts were prepared normally and imaged by TIRF microscopy to verify their assembly. All solutions prepared for SEM were passed a 0.2 μm syringe filter. Reaction buffer was removed, and samples were fixed with 2% glutaraldehyde in 0.05M sodium cacodylate (NaCaco) buffer for 30 minutes. Fixed samples were rinsed twice for 5 min with 0.05M NaCaco buffer before incubated with 0.5% osmium tetroxide for 15 min. After rinsing twice with NaCaco buffer for 15 min, samples were treated with 1% tannic acid for 15 min, followed by three additional 15 min rinses. Samples were then treated a final time with osmium tetroxide for 15 min and rinsed twice for 5 min with NaCaco buffer. Samples were then dehydrated in series with ethanol (30%, EtOH 5 min, 50% EtOH 5 min, 75% EtOH 5 min, 100% EtOH twice for 5 min, once more for 10 min). Samples were dried for 15 min with CO_2_ using a Samdri-795 critical point dryer (Tousimis Research Corporation) before sputter coating (Cressington Sputter Coater 208HR, model 8000-220, Ted Pella). Samples were coated with a 4 nm layer of gold/palladium (60:40) alloy before imaging on a Zeiss Supra 25 Field Emission scanning electron microscope.

### *Ashbya gossypii* culturing and imaging

*A. gossypii* (Ag*CDC11a*-GFP:GEN leu2Δ thr4Δ) was grown from spores in full medium at 30°C for 16 hours before harvesting mycelia. Full media was washed out of solution using low fluorescence media. Mycelial cells were mounted onto low fluorescence medium pads solidified with 2% agarose. Images were collected with a TI-LA-HTIRF (Nikon) microscope (Nikon Ti-82 stage) using a 100x Plan Apo 1.49 NA oil objective and a Prime 95B cMOS camera (Photometrics).

### Constructing a physical model to simulate isodesmic septin polymerization

Brownian dynamics simulations were run in HOOMD-Blue (Anderson et al., 2020) augmented with a custom C++ plugin that implemented dynamic bond and angle formation and deletion, and ghosting reactions. The plugin was adapted from a plugin used in a previous work to model epoxy curing (Thomas et al., 2018). Simulations were run on GPU nodes on the Longleaf computer cluster at UNC-Chapel Hill.

### Model specifications

Octamers were modeled as circles constrained to a two-dimensional plane with periodic boundary conditions. We initially considered the octamers to be spheres that would be free to rotate in three dimensions. However, it seems likely that the octamer rods bind to the SLB with a highly restricted interface, meaning that the bound octamers have no rotation about their long axis. In this case the only rotation is that of the long axis within the plane of the SLB, reducing the octamer to a circle.

We used Brownian dynamics to simulate their thermal diffusion:

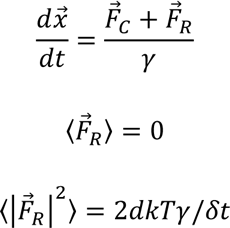

where 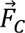 is the implicit force on the particle (octamer) from all potentials (e.g. excluded volume interactions, harmonic bonds, angle potentials), 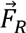 is a uniform random force, *γ* is the drag coefficient, *d* is the dimensionality of the system, and *kT* is the Boltzmann coefficient. In our system, *d* = 2, and we set the drag coefficient and the Boltzmann coefficient equal to 1, scaling other quantities accordingly. Our time step, *dt*, was set to 1e-4 (arbitrary time units).

Particles experienced excluded volume interactions (actually excluded area in the two-dimensional model) which were implemented using a Lennard-Jones pair potential:

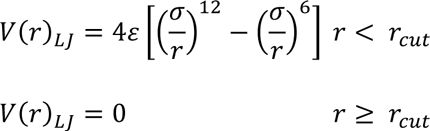

with *ε* = 5, *σ* = 1, and **r*_cut_* = 2^1⁄6^.

To model polymerization/fragmentation and membrane binding/unbinding reactions, we supplemented the Brownian dynamics framework with a custom plugin. This plugin evaluated and updated the states of all particles in the system every 1e4 time steps; we will refer to this evaluation as the update step. To model membrane binding and unbinding, we implemented a “ghosting” reaction. Monomers on the membrane ghost with a probability *k_off_*, which is evaluated for each monomer (unbound particle) during each update step. A ghost monomer simulates a monomer that has unbound from the membrane and is diffusing through solution. Therefore, “ghosts” no longer experience excluded volume interactions (*ε* = 0 in all of its pairwise Lennard-Jones interactions), diffuse much faster (*γ* = 0.01 so diffusion rate is 100x that of a membrane bound particle), and are ineligible for polymerization reactions with other particles. Incorporating “ghosting” into the model is computationally less expensive than “destroying” octamers that unbind the membrane and “respawning” them at random location. During an update step, if a ghost particle is not overlapping with any membrane-bound particles, it can bind the membrane with a probability *k_on_*, and the corresponding parameters are set back to their default values. *k_on_* and *k_off_* were parameterized as relative probabilities proportional to the average “spaced explored” in two-dimensions (i.e. the average displacement of a membrane-bound octamer by diffusion in two-dimensions). We set the distance unit to be the length of an octamer (i.e., diameter of the sphere), 32 nm. The energy unit is set as *k*T at room temperature, 2.479 kJ/mol. The mass is set to be the mass of an octamer, 360 kD. Note we used a diffusion coefficient of ∼1.8 *μm*^2^/*s*, which is about 10 times the measured diffusion rate. The measured octamer dissociate rate from the membrane or *k_off_* = 1.42 *s*^−1^. In the model, the “space explored” by an octamer on the membrane translates into an average octamer “exploring” a space of 2228 square simulation units during an average membrane-bound lifetime. Thus, in the model we set *k_off_* probability to be 4.5e-4 (i.e. 1/2228). Speeding up diffusion reduced the simulation time to reach steady state but did not otherwise affect steady state properties. To obtain an octamer association rate for the model, we measured the ratio of membrane-bound Cdc11α6-capped octamers relative to the concentration in solution at steady state. This ratio suggested that 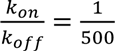, thus the membrane *k_on_* probability in the model is 9e-7.

We modeled polymerization reactions according to several categories of interacting particles. Only particles bound to the membrane can polymerize. Additionally, particles are divalent spheres and can only bind a maximum of two other particles. If two monomers touch (their center-to-center distance is less than or equal to a particle diameter times 2^1⁄6^), they bond with a probability *P_bond_*= (2*θ*/360)^2^, where *θ* is a tolerated binding angle. Since spherical particles represent non-polar rod-like septin octamers, the probability of binding simulates freely rotating rods that polymerize only where their ends are aligned within a specified angle tolerance. We set *θ* = 10^∘^ which is extracted from the measured septin filament persistence length (L_P_; i.e., flexibility). A freely diffusing octamer anneals encountering an existing filament anneals with a probability *P_bond_* = 2*θ*/360, since the octamer at the filament end is assumed to be aligned with the two end subunits of the filament whereas the octamer is freely rotating. Two filaments in close proximity will anneal if both of their terminal subunits are within *θ*. The following harmonic bond potential is added between the two polymerized octamers:

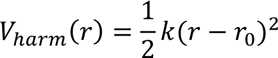

where *k* is the potential constant (set to 40), *r* is the distance between the pair of particles being considered, and *r*_0_ is the rest bond length (set to 2^1⁄6^ times the diameter of a particle).

We implemented persistence length by adding a cosine squared angle potential between every consecutive triplet of octamers within filaments as they polymerize:

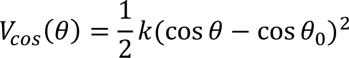

where *k* is the potential constant, *θ* is the angle between the triplet of particles, and *θ*_0_ is the rest angle (set to 180^∘^). The value of *k* sets the persistence length of a filament according to the following relation (Zhang et al., 2019):

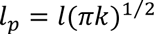

where **l*_p_* is the persistence length, *l* is the monomer diameter, and *k* is the potential constant in *V_cos_*. Specification of a persistence length allows for a specification of *k* in the simulation. Note that persistence length also likely emerges from bending between subunit interactions with the octamer, thus modeling filament persistence length by allowing only bending between octamers is an approximation (Gittes et al., 1993). Given a persistence length, the average angle between monomers in a filament is given by:

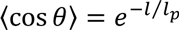

The measured persistence length and a particle diameter of 32 nm (length of an octamer) yields an average bond angle of 10 degrees. However, when altering persistence length in simulations parameter sweeps, the tolerated bond angle was maintained at 10 degrees to isolate the effect of a change in persistence length.

Isodesmic filaments fragment anywhere along their length with equal probability. This was modeled with a fragmentation probability *P_frag_* = *e^-E/kT^*, where *E* is the binding energy between septins. When bonds are broken, the harmonic bond potential between the pair of participating particles is removed, and all corresponding angle potentials are removed.

## Acknowledgements

We thank Harold Erickson, Masayaki Onishi, Daniel Lew, Ehssan Nazockdast, and Wenzheng Shi for comments on the manuscript, we thank members of the A.S.G. laboratory for stimulating discussions on this research project, and we thank Victoria Madden and Kristen White of the UNC electron microscopy core facility for help with SEM.

This work was supported by NSF Grants 1615138 and 20216022 and the Howard Hughes Medical Institute Faculty Scholars program awarded to A.S.G. B.L.W. was supported by the NIH Training Grant 2T32AI052080-16. K.S.C. was supported in part by a grant from the National Institute of General Medical Sciences under award T32 GM119999.

## Notes

### Competing Interest Statement

The authors have declared no competing interest.

